# Spatially resolved transcriptomic profiling of degraded and challenging fresh frozen samples

**DOI:** 10.1101/2022.09.13.507728

**Authors:** Reza Mirzazadeh, Zaneta Andrusivova, Ludvig Larsson, Phillip T. Newton, Leire Alonso Galicia, Xesús M. Abalo, Mahtab Avijgan, Linda Kvastad, Alexandre Denadai-Souza, Nathalie Stakenborg, Alexandra B. Firsova, Alia Shamikh, Aleksandra Jurek, Niklas Schultz, Monica Nistér, Christos Samakovlis, Guy Boeckxstaens, Joakim Lundeberg

## Abstract

Spatially resolved transcriptomics (SRT) has enabled precise genome-wide mRNA expression profiling within tissue sections. The performance of unbiased SRT methods targeting the polyA tail of mRNA, relies on the availability of specimens with high RNA quality. Moreover, the high cost of currently available SRT assays requires a careful sample screening process to increase the chance of obtaining high-quality data. Indeed, the upfront analysis of RNA quality can show considerable variability due to sample handling, storage, and/or intrinsic factors. We present RNA-Rescue Spatial Transcriptomics (RRST), an SRT workflow designed to improve mRNA recovery from fresh frozen (FF) specimens with moderate to low RNA quality. First, we provide a benchmark of RRST against the standard Visium spatial gene expression protocol on high RNA quality samples represented by mouse brain and prostate cancer samples. Then, we demonstrate the RRST protocol on tissue sections collected from 5 challenging tissue types, including: human lung, colon, small intestine, pediatric brain tumor, and mouse bone/cartilage. In total, we analyzed 52 tissue sections and our results demonstrate that RRST is a versatile, powerful, and reproducible protocol for FF specimens of different qualities and origins.

## Introduction

Spatially resolved transcriptomics (SRT) is a set of technologies used to chart genome-wide mRNA expression within tissue sections, and it has become widely used in genomics research in the past decade ^1–3^. SRT has opened up new possibilities to explore the spatial architecture of cells and their interactions in the tissue context, exemplified by works in neuroscience ^4^, developmental biology ^5^, and disease ^6,7^.

The first report of such an SRT method for high throughput spatial mRNA profiling was published in 2016 ^8^. This work paved the way for unbiased capturing of whole transcriptomes from tissue sections. The underlying principle of this technology is a dense grid of spatially barcoded oligo(dT) probes printed on a microscope glass slide, which can be used to capture the polyA tails of mRNA molecules from a tissue section, thus facilitating spatially resolved gene expression profiling. The tissue section is also stained and imaged with a microscope, which makes it possible to combine gene expression profiling with histology. Currently, the most broadly used SRT platform is Visium (10x Genomics) ^3^, an updated version of the same principles presented by Ståhl *et al*.,^8^ currently with 5000 barcoded spots, each with a diameter of 55μm (see 10x Genomics webpage ^9^).

The Visium protocol is optimized for fresh frozen (FF) tissue specimens and recommends a RIN (RNA Integrity Number) score higher than or equal to 7. RIN is a critical metric to assess the quality and level of RNA degradation before starting an SRT experiment ^10,11^. FF samples are the preferred choice for unbiased polyA-based SRT technologies due their high preservation of polyadenylated transcripts. However, a major limitation with polyA-based SRT is its reduced ability to process degraded samples. In spite of the widespread use of Visium for FF samples, there is a need for a method that works well on samples with low RNA quality. Recently, 10x Genomics introduced a new chemistry for Formalin-Fixed Paraffin-Embedded (FFPE) samples. In FFPE samples, it is well documented that RNA molecules are fragmented, where the degradation often affects the polyA tails of the RNA ^12,13^. To overcome the aforementioned issue, the FFPE SRT approach relies on a gene-panel to target and capture protein-coding regions of the transcriptome instead of targeting the polyA tails.

Based on this recent development, we propose a strategy for spatial analysis of FF tissue specimens with moderate/low RIN scores, that we name RNA-Rescue Spatial Transcriptomics (RRST). This protocol makes use of the same targeted gene-panel that was designed for FFPE material with additional modifications to work on FF tissues, including a gentle formalin fixation step and a baking step to improve tissue adherence to the slide surface. We demonstrate the capabilities of our RRST method by profiling the tissue transcriptomes of a variety of biological specimens, and comparing the results with data generated by the standard Visium protocol.

## Results

### RRST implementation in fresh frozen tissue sections

We attempted to make the Visium SRT technology compatible for analysis of degraded FF samples by introducing specific modifications to the commercially available Visium FFPE protocol. In the original FFPE protocol, tissue sections are first deparaffinized through a series of washes with xylene/ethanol. Then, the tissue sections are stained with hematoxylin-eosin and de-crosslinked in the Tris-EDTA (TE) buffer at 70 °C for an hour. The sections are then incubated with probe sets that hybridize in pairs to each transcript, targeting approximately 19K proteincoding genes. Upon correct probe hybridization to mRNA transcripts, each pair is ligated to one another and captured by oligo(dT) probes attached to the surface of the glass slide, where spatial barcodes are introduced through a cDNA synthesis step. The cDNA molecules, which now hold information about the target transcript and its spatial location, are released from the slide surface for final library preparation and sequencing.

It should be noted that FFPE specimens are usually heavily crosslinked due to a prolonged formalin fixation process, and thus crosslink reversal is a critical step to access the RNA molecules within tissue sections. This reversal is done through long incubation at high pH and temperature. In relation to this, fairly harsh treatment of tissue sections and long incubation at high temperature during Visium FFPE protocol increase the chance of tissue detachment in initial steps of section processing, which may result in low quality/failed data generation (**Supplementary Fig. 1, Supplementary Video 1**). In relation to this, our RRST protocol is designed to account for all of these factors which is a key to successful spatial gene expression profiling of FF tissue samples. In the RRST protocol, FF tissue sections are fixed with formalin, instead of methanol, for 10 min at room temperature, followed by a baking step of 20 minutes at 37°C, which we found necessary in order to improve tissue section adhesion to the Visium slides. Since RRST protocol introduces short formalin fixation, we speculated that the long decrosslinking incubation step at high pH and temperature used in FFPE protocol may potentially lead to RNA degradation in the FF samples. Hence, we removed the cross-linking reversal step in our protocol, which in addition shortens the duration of protocol by an hour. A detailed workflow of the RRST is depicted in **Supplementary Fig. 2** and a step-by-step protocol can be found in the Methods section.

### Performance of RRST in high quality FF samples

We first set out a test to evaluate how well the RRST protocol performs on two FF samples with high RIN values: a mouse brain sample (RIN 8.8) and a human prostate tumor specimen (RIN 10) (**Supplementary Table 1**). The mouse brain has become the sample of choice to benchmark SRT technologies because of its well-defined anatomical structures, which have been characterized in detail based on histology and spatial gene expression ^14^. We performed both RRST and standard Visium on these samples and found that RRST can robustly profile the tissue transcriptome with approximately 2-fold increase in the number of detected genes per spot compared to the standard protocol **(Fig. 1b, c)**. Moreover, in both the mouse brain and prostate tumor samples, we observed a high concordance (Pearson R = 0.82, p < 2.2e-16 and R = 0.76, p < 2.2e-16) between the aggregated gene counts across the two datasets, excluding genes that were not targeted by the RRST panel (**Fig. 1d**). This indicates that the data obtained with the RRST approach display a high similarity with the data obtained with the standard Visium protocol. However, the probe panel used for RRST excludes certain transcripts, such as those transcribed from mitochondrial genes, ribosomal protein coding genes or ncRNAs (**Supplementary Fig. 3**). With the exception of these three RNA types, the majority of detected transcripts come from protein coding genes and are detected with both methods (**Supplementary Fig. 3**), although at drastically different UMI counts and detection rates (**Fig. 1d**). For the majority of transcripts, RRST protocol appears to exhibit a higher capture efficiency.

**Fig.1:**
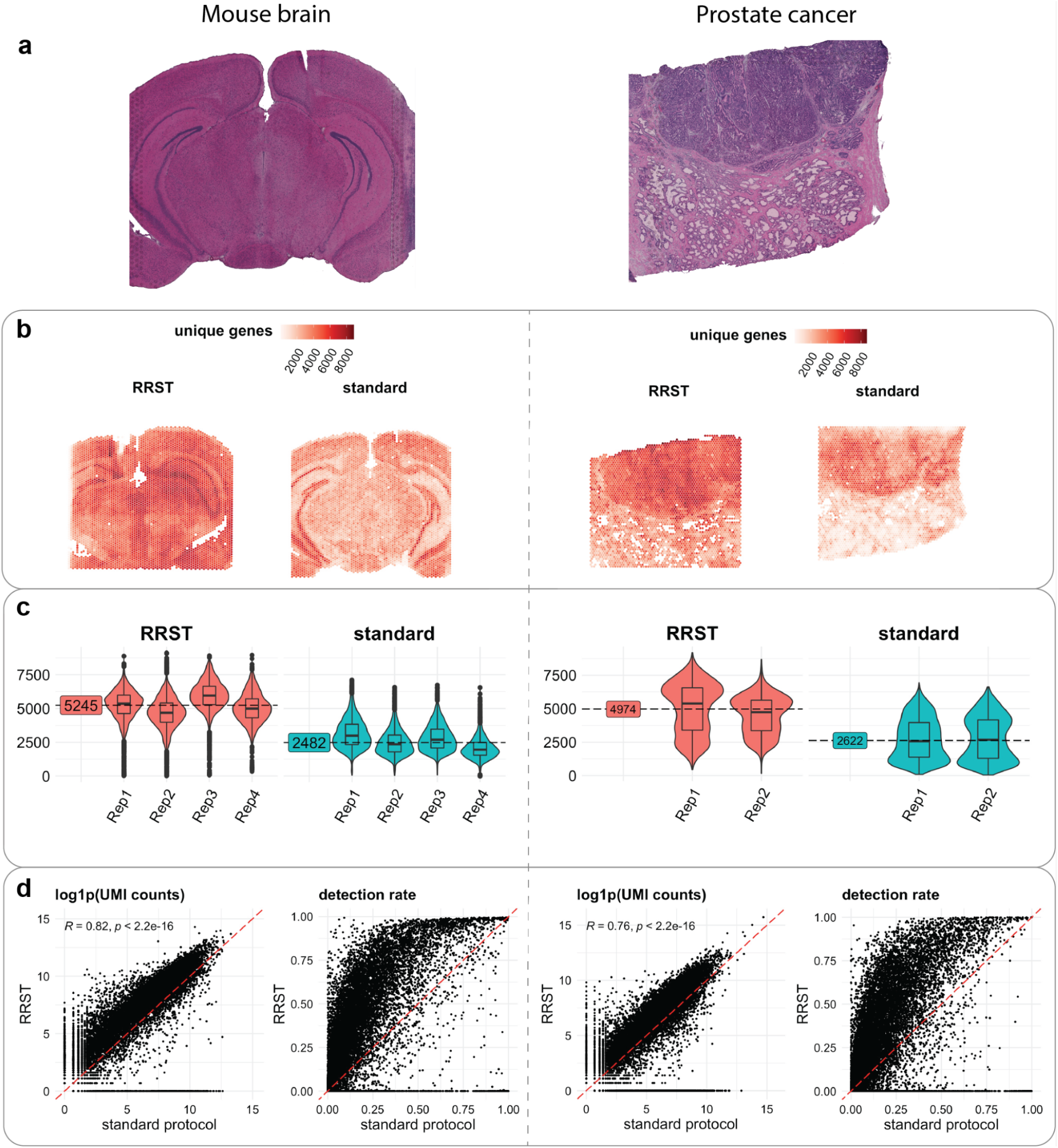
Comparison of RRST and Visium on mouse brain and prostate tumor samples. **a)** H&E images of a representative tissue section from mouse brain (left) and prostate cancer (right) **b)** Spatial distribution of unique genes in two representative tissue sections for each tissue type, one processed with the RRST protocol and one processed with the standard Visium protocol. **c)** Distributions of unique genes per spot visualized as violin plots colored by experimental protocol for mouse brain (n = 8) and prostate cancer (n = 4) data. The median number of unique genes is highlighted for each group (sample type and protocol) next to the violin plots. **d)** gene-gene scatter plots between RRST data (y-axis) and standard Visium data (x-axis) of log1p-transformed UMI counts or detection rates using the data shown in **b**. The red dashed line highlights a 1-to-1 relationship. For the log1p-transformed UMI counts scatter plot, only genes targeted by the probe panel were included. The detection rate for a gene is defined as the proportion of spots with detected UMI counts.

### RRST recovers spatial transcriptomics data from challenging FF samples

There is a growing number of studies using the standard Visium platform for FF tissues to address biological questions ^15^. The assessment of RNA quality through RIN measurement (RIN ≥ 7) is suggested as an important criterion to define the quality of tissues for successful spatial gene expression profiling. In our own experience, some tissue types are more challenging to retrieve good/high quality Visium data from. There could be several factors contributing to low/moderate RNA quality, such as intrinsic biological characteristics of the tissue, rapid RNA degradation upon surgical procedure or sensitivity to freezing/thawing during tissue sectioning. Hence, we aimed to apply RRST to some challenging tissue types that are known to perform poorly using the standard 3’ capture Visium platform (**Supplementary Figs. 4, 5**).

#### Adult human lung tissue

To date, spatial transcriptome profiling of human lung tissue has rarely been investigated ^16^. Based on our own experience, FF mouse and human lung tissue samples are highly challenging to process with the standard Visium protocol. Therefore, we tested the performance of RRST in FF healthy adult human lung samples (**Fig. 2a, Supplementary Fig. 6a**) retrieved from two patients (LNG1, RIN 6.8 and LNG2, RIN 7.1, **Supplementary Table 1**), where the 3’ capture protocol performed poorly. Our RRST method detected roughly a 2-fold and 10-fold increase in the number of detected genes per spot in these two patients respectively, indicating the robustness and power of RRST to profile spatial gene expression in challenging tissue types (**Fig. 2b, c and Supplementary Fig. 6b, c**). As a quality control step, spots with few unique genes detected are commonly discarded based on an empirical cutoff threshold, where thresholds between 500-1000 unique genes are common. Here we used a softer cutoff threshold of 300 unique genes to include as many spots as possible from both conditions **(Fig. 2d, Supplementary Fig. 6d**). Even with this soft cutoff threshold, ~21%-80% of the spots were discarded for the standard Visium data, while only 1.3%-2.3% of spots for the RRST data from the same tissue blocks (**Supplementary Fig. 7**). For downstream analysis, we first focused on one of the patients (LNG1). After dimensionality reduction and clustering using the same parameter settings for the two data types, we were able to detect 11 clusters in the RRST and 9 clusters in the standard Visium data (**Fig. 2e, f**). Notably, marker detection by Differential Expression Analysis (DEA) highlighted distinct marker genes for each of the 11 RRST lung clusters whereas clusters 0, 1 and 2 in the standard Visium lung data were difficult to distinguish from each other (**Fig. 2g**). Moreover, cluster 4 in the standard Visium lung data displayed differential expression of mitochondrial transcripts, which is indicative of low quality transcriptomic profiles ^17^. Interestingly, some of the top markers detected for cluster 8 (airway epithelium) in the standard Visium lung data were ncRNAs (*LINC00326, ACBD3-AS1* and *AC023300.2*), which RRST does not detect. We characterized the RRST clusters based on the top markers detected and the spatial localization (**Fig. 2f, g and Supplementary Fig. 8a**). Next, we inspected the expression of top markers detected in four selected clusters in the RRST data: airway epithelium, megakaryocyte-enriched, smooth muscle and glands. We found that the expression levels of these markers were consistently higher in the RRST data, in line with the higher data quality and complexity (**Supplementary Fig. 8b**). In addition, we applied the same analysis workflow on the second patient sample (LNG2) observing similar trends of higher RRST data quality in contrast to standard Visum protocol (**Supplementary Fig. 6**). Together, these results indicate that RRST provides increased power to profile and detect transcripts in lung tissue specimens.

**Fig.2:**
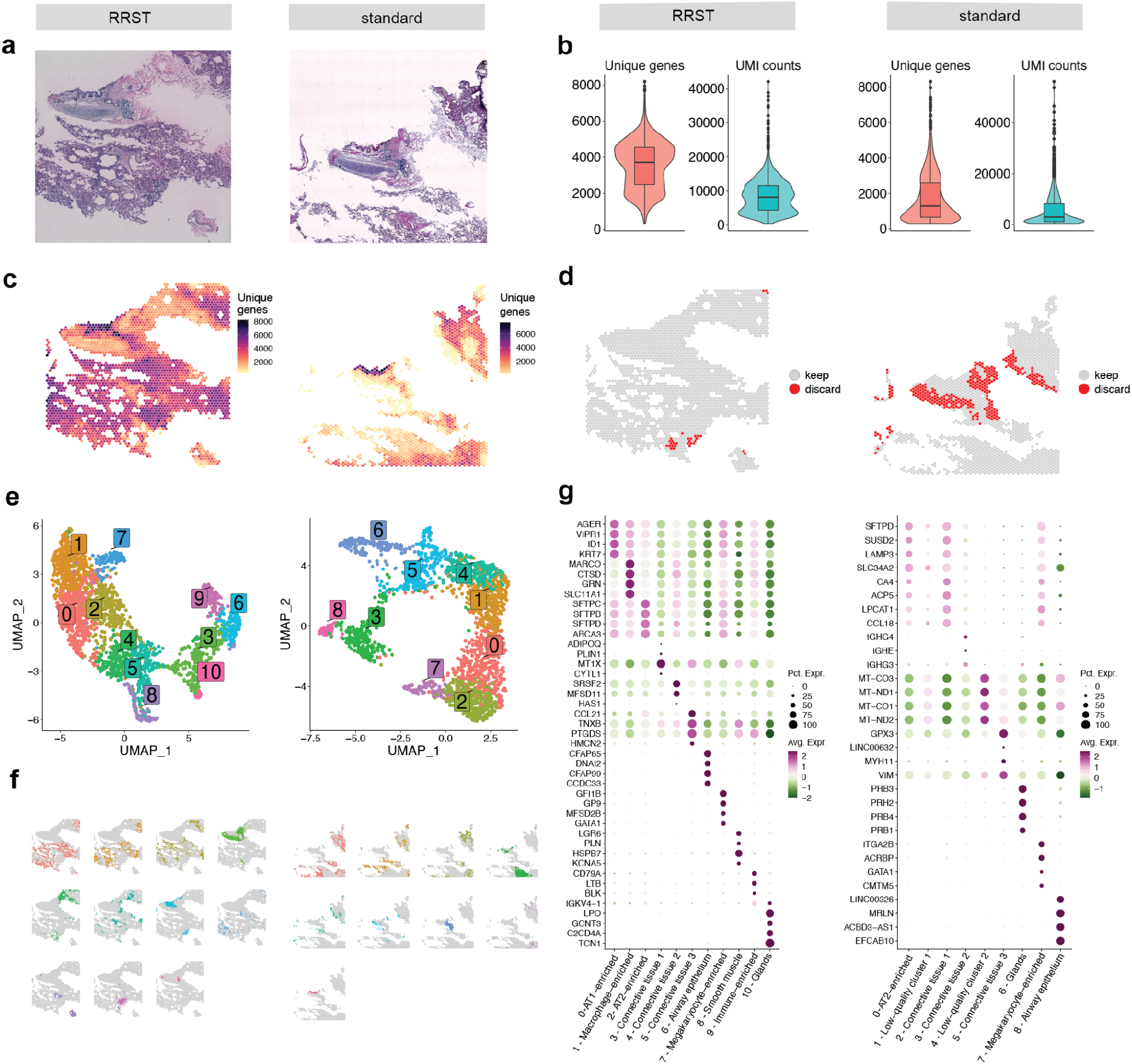
RRST and standard Visium applied to human adult lung tissue. Each subplot shows the RRST data on the left side and the standard Visium data on the right side. **a)** H&E images of two representative tissue sections collected from the same tissue block. **b)** Violin plots showing the distribution of unique genes and UMI counts. **c)** Unique genes per spot mapped on tissue coordinates. **d)** Spatial visualization showing what spots were discarded due to low quality (less than 300 unique genes detected). **e)** UMAP embedding of adult lung data colored by clusters detected by unsupervised graph-based clustering (louvain). **f)** Split view of clusters (same as in **e**) mapped on tissue coordinates. **g)** Dot plots of the top marker genes for each cluster. Each cluster was annotated based on its spatial localization in the tissue and expression of canonical marker genes.

#### Adult human colon tissue

Next, we investigated if the RRST protocol can be used to obtain spatial gene expression data from samples for which the standard Visium failed. For this purpose, we investigated FF adult human colon samples collected for the Gut Cell Atlas consortium. After extensive efforts to generate good-quality data from these tissue blocks, we could conclude that the intestinal epithelial tissues are particularly susceptible to mRNA degradation and consequently difficult to process with the standard Visium protocol. It is of note that the gut is a highly delicate tissue that is filled with digestive enzymes and a microbiome of varying quality and quantity, which in turn can lead to a rapid degradation of RNA ^18^. We processed colon tissue sections obtained from two patients (**Fig. 3a**) with moderate RNA integrity (RIN of 4.5 and 5.1, **Supplementary Table 1**). To assess whether mRNA degradation differs between tissue types, we manually annotated the data into three major regions: mucosa, submucosa and muscularis (**Fig. 3a**). Notably, in the standard Visium data, we observed low numbers of unique genes and UMI counts in the cell dense epithelial layer (mucosa) while we could still recover decent numbers of unique genes and UMIs counts in the muscularis (**Fig. 3b-d**). This observation was in line with what has been reported previously in literature, where it was shown that mRNA degrades more rapidly in the intestinal epithelium compared to the intestinal muscle tissue ^18^. However, with the RRST protocol, we were able to recover good-quality data both from the mucosa and submucosa in tissue sections collected from the same OCT block (**Fig. 3b-d**). The RRST method generated more even data coverage across different tissue regions (**Fig. 3b**), indicating that the method is able to mitigate the effects of tissuespecific degradation. To demonstrate the effect of tissue-specific degradation, we investigated expression in the mucosa of 11 intestinal epithelial markers (**Fig. 3e**) selected from the Gut Cell Atlas ^19^. These results show that the RRST data provided higher detection rates and more even expression values, thus indicating that the method can be used to profile regions with degraded mRNA.

**Fig.3:**
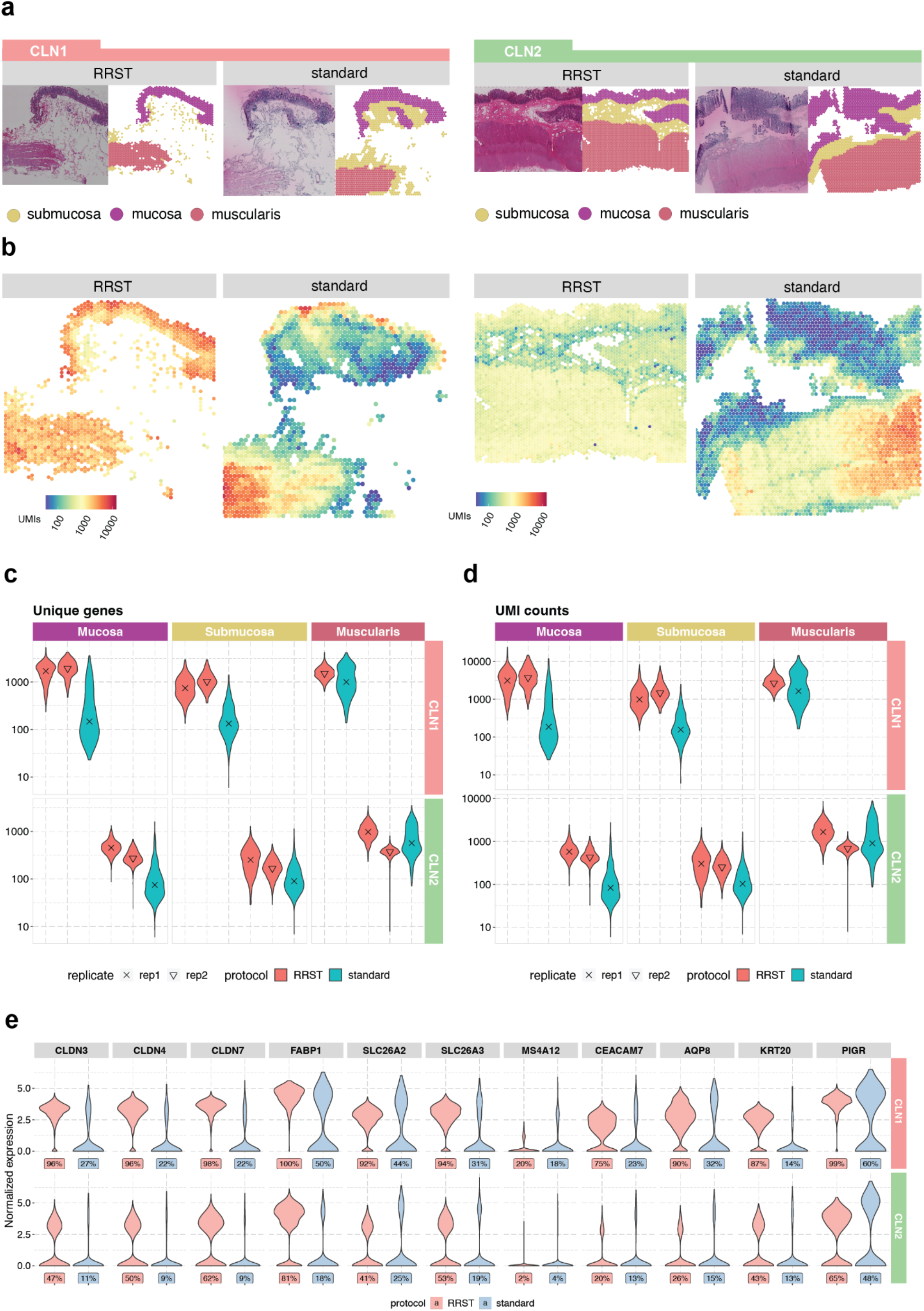
Comparison of data quality in RRST and standard Visium datasets generated from adult human colon tissues. **a)** Representative H&E images and annotated regions for two patient samples, processed by either RRST (n = 4) or standard Visium (n = 2). The spots in each tissue section were labeled into three categories: mucosa, submucosa and muscularis. **b)** Distribution of UMI counts in the tissue sections shown in **a**. The color scale represents log10-transformed counts. **c)** Distribution of unique genes per spot in the three annotated regions (mucosa, submucosa and muscularis) visualized as violin plots, for all tissue sections. The y-axis shows log10-transformed counts. **d)** Distribution of UMI counts per spot in the three annotated regions (mucosa, submucosa and muscularis) visualized as violin plots. The y-axis shows log10-transformed counts. **e)** Expression of 11 epithelial markers in the mucosa for the two adult colon samples visualized as violin plots. A comparison between the two protocols is shown for each gene and the corresponding detection rate is highlighted below each violin plot. The detection rate is defined as the percentage of spots (in the mucosa) where the gene is detected.

#### Adult human small intestine tissue

The mRNA quality of FF tissue blocks depends on a number of different factors such as sample collection, handling and storage ^20^. To estimate the overall quality of a specimen, it can be useful to measure RIN and/or DV200. However, for certain sample types, we have observed that mRNA can degrade rapidly even when they are properly stored in freezers, which in turn means that quality measurements become less reliable over time. One such sample, where we could observe a rapid degradation, was a FF OCT-embedded tissue specimen from an adult human small intestine (Ileum) obtained from the Gut Cell Atlas project. Approximately 1 month after sample collection, we processed four tissue sections from the FF OCT block using the standard Visium protocol which generated high-quality data from all tissue regions: mucosa, Tertiary Lymphoid Tissue (TLS), submucosa, muscularis and serosa (**Fig. 4a-b, Supplementary Fig. 9**). Surprisingly, when we repeated the experiment using the same tissue block six months later (8 tissue sections), we observed an almost complete loss of gene expression data in the mucosal/submucosal layers, while the data in the muscularis remained stable and comparable to the first experiment **(Fig. 4b)**. These results reiterate what we observed in the adult human colon tissue, that mRNA degradation can vary in different tissue types and even within the same section, which cannot be assessed by bulk RIN quality check prior to the SRT assay. Moreover, it became clearer that the main challenge with running Visium on intestinal lower GI tract epithelial tissues is the rapid mRNA degradation. To test whether we could use our RRST method to recover high-quality data from the same block, we processed two tissue sections from the same OCT block approximately two years after sample collection (RIN 7.8, **Supplementary Table 1**). As indicated by the number of unique genes, we were able to detect higher numbers in the mucosa and submucosa compared to the second attempt, albeit with lower numbers than the initial experiment conducted two years earlier (**Fig. 4b**).

**Fig.4:**
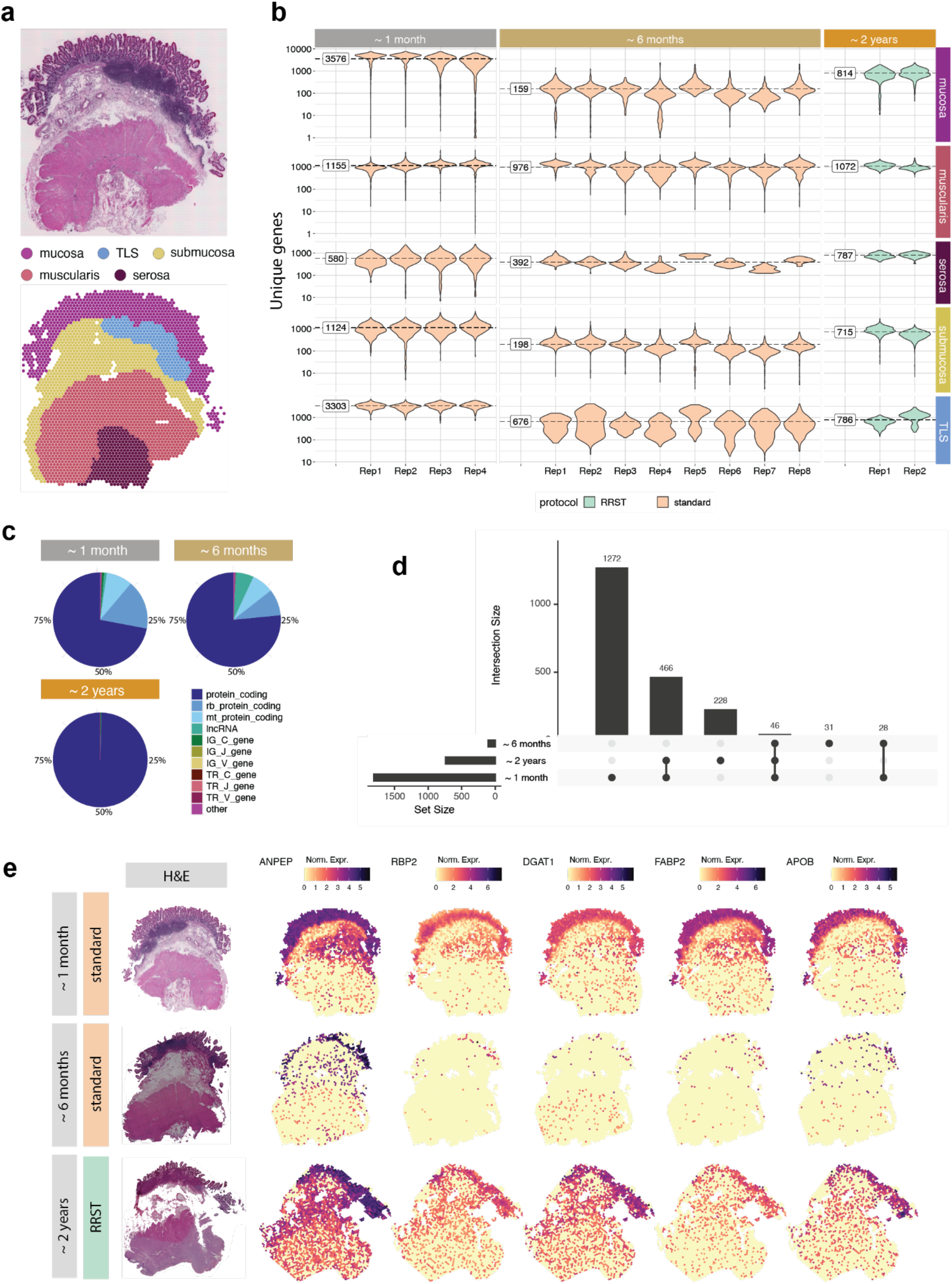
Comparison between RRST and standard Visium on an adult human small intestine sample over time. **a)** H&E image (top) and spots colored by five major tissue regions (bottom): mucosa, TLS, submucosa, muscularis and serosa. TLS, Tertiary Lymphoid Tissue. **b)** Overview of data quality in the five annotated tissue regions over time, visualized by violin plots of the number of unique genes per spot. The time points represent the approximate storage time after sample collection: ~ 1 month, ~ 6 months and ~ 2 years. Replicates obtained for each time point are shown on the x axis. The fill color of the violin plots indicates the protocol used. For each time point, labels on the left side of the violin plots represent the average over all replicates. **c)** RNA biotype content for the three datasets visualized as a pie chart. Proportions represent the UMI counts detected for each biotype. The targeted RRST data include protein coding, immunoglobulin and T-cell receptor transcripts. **d)** Upset plot highlighting the number of differentially expressed genes (DEGs) detected in the mucosa in each of the three time points (average log2-fold change < 0.25, adjusted p-value < 0.01). Horizontal bars on the bottom left represent the total number of DEGs for each time point. Vertical bars on the top represent the intersection sizes for DEGs that were detected uniquely or in one time point or in a combination of multiple time points. Intersections are highlighted under the vertical bars. **e)** Spatial visualization of five enterocyte markers. Each row represents one selected tissue section from each time point with their corresponding H&E image in the leftmost column. Spot colors represent normalized gene expression.

Notably, the second attempt with the standard Visium method (~ 6 months after sample collection) resulted in an average of 159 unique genes per spot in the mucosa, whereas the RRST data (~ 2 years after sample collection) resulted in an average of 814 unique genes per spot in the mucosa (**Fig. 4b**). Moreover, a large fraction of the expression data obtained with standard Visium comes from mitochondrial transcripts, ribosomal protein coding transcripts or lncRNA, which are commonly filtered out prior to downstream analysis, whereas RRST only targets protein coding genes (**Fig. 4c**). Next, we looked closer at the mucosa region to determine how the difference in quality affects the ability to detect differentially expressed genes. For each time point, we ran differential expression analysis between the mucosa and the remaining tissue regions. We found that while most of the genes (1272) were only detected in the initial dataset, 466 genes were detected both in the initial dataset and the RRST dataset and 228 genes were uniquely detected in the RRST dataset (**Fig. 4d**). On the other hand, only 31 differentially expressed genes could be detected in the standard Visium dataset produced 6 months after tissue collection. Next, we took five enterocyte markers from the Gut Cell Atlas and visualized their expression across the tissue sections in the three datasets. These markers were clearly visible in the mucosa in the first dataset and the RRST dataset but not in the second dataset (**Fig. 4e**). Based on these results, we speculate that rapidly gut epithelial tissues contain high amounts of RNAses and therefore repetitive freeze/thaw cycles and long-term storage lead to mRNA degradation, hence our RRST approach can help overcome these effects. Overall, these results demonstrate that our RRST protocol can be used in FF samples with low/moderate RNA integrity and to recover data from FF tissue blocks that have been stored for long periods of time.

### RRST revives spatial transcriptome profiles of precious clinical samples

Spatial gene-expression profiling of clinical samples can enable discoveries required to develop new strategies for early diagnosis and individualized therapies at molecular levels ^21^. Treatment of pediatric brain tumors is continually being improved upon; however, there is a great need for new treatment options. Due to the limited amount of tissue available for research, there is usually not enough material for tissue optimization and RIN measurement to assess whether the sample quality is sufficient for the standard 3’ capture protocol. In order to investigate how RRST performs in such precious clinical samples, we processed two pediatric brain tumor specimens (RIN 7.0 and 7.1, **Supplementary Table 1**) from which we had previously failed to generate data using the standard 3’ polyA capture protocol. In contrast to previous samples described in this study, the pediatric brain tumor samples passed the recommended RNA quality threshold for the standard Visium assay. We speculate that the underlying reason for why these experiments failed was due to either tissue detachment or inefficient permeabilization of the tissue. By applying RRST protocol to these samples, we could reach a 12 to 100-folds increase in the number of detected genes per spot (**Fig. 5a, b**). This suggests that the RRST approach is less sensitive to changes in tissue composition compared with the standard Visium protocol.

**Fig.5:**
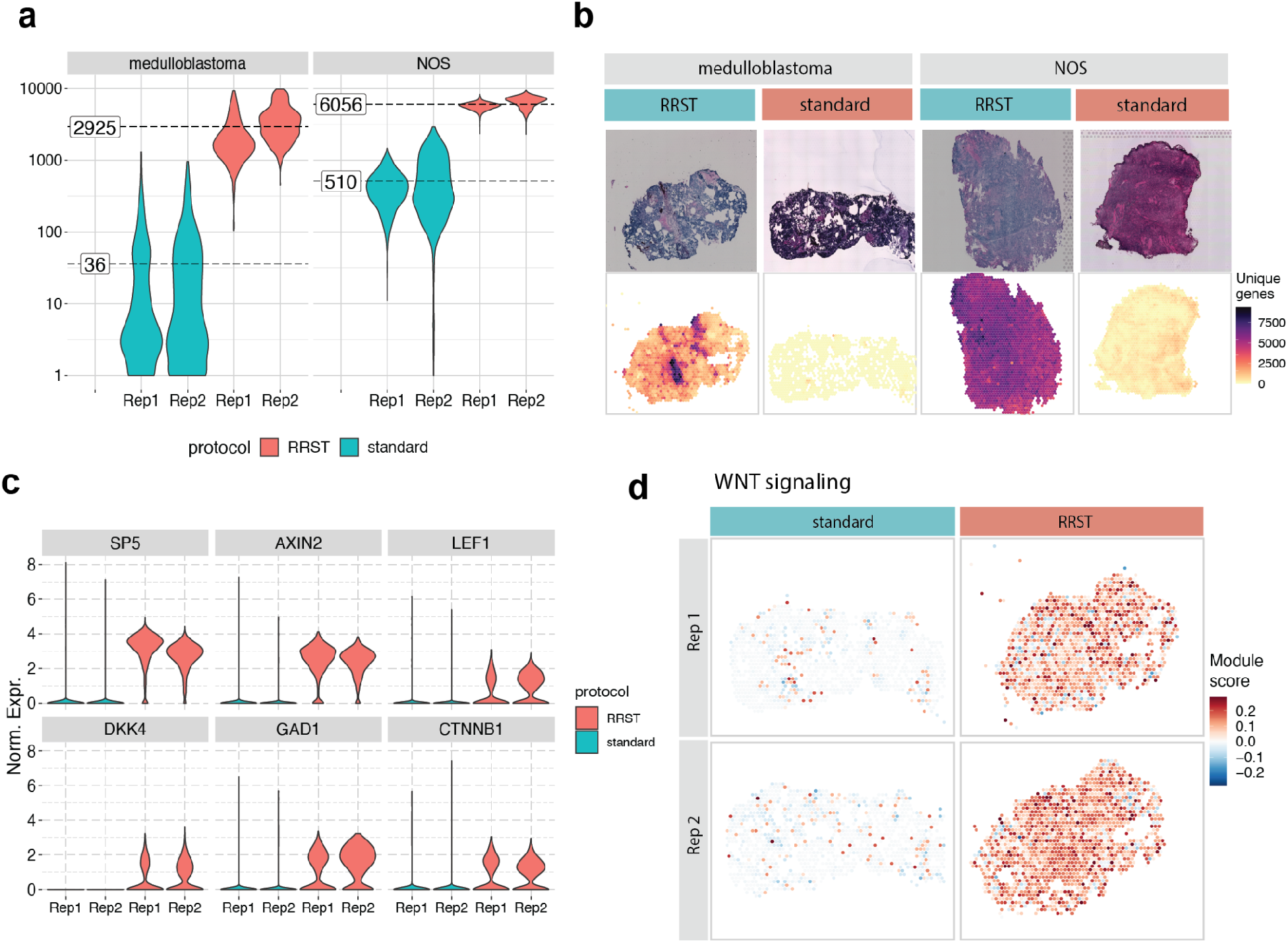
Comparison between standard Visium and RRST protocols in eight pediatric brain tumor tissue sections. **a)** Violin plots showing the number of unique genes per spot in all eight tissue sections (medulloblastoma n=4, NOS n=4). The fill color represents the protocol used to generate the data. The average number of unique genes for each sample and protocol are highlighted by dashed lines. Note that the y-axis shows log10-transformed values. **b)** H&E images (top row) and the number of unique genes per spot (bottom row) shown for four representative tissue sections. **c)** Violin plots showing the normalized expression of 6 marker genes related to WNT-signaling across the four medulloblastoma tissue sections. The fill color represents the protocol used to generate the data. Rep1, replicate 1; Rep2, replicate 2. Norm. Expr., normalized gene expression. **d)** Spatial visualization of WNT--signaling module scores in the WNT medulloblastoma samples.

Based on the low data quality of the standard Visium data, we were first discouraged to proceed with data analysis. However, with the RRST data, we could assess how the difference in quality affects characterization of these tumors. For this purpose, we focused on the medulloblastoma sample which was classified as a WNT subtype, characterized by activation of the WNT signaling pathway ^22^. The medulloblastoma tissue sections were annotated by a pathologist, showing that most of the tissue sections were composed of tumor cells (**Supplementary Fig. 10**). To compare the data quality of RRST and standard Visium datasets obtained from the pediatric brain tumor samples, we examined the expression of WNT-signaling genes, including *AXIN2, DKK4, LEF1 and CTNNB1* and 2 known targets of the WNT pathway *SP5, GAD1* ^23^,24. We were able to detect these marker genes in the RRST dataset, but not in the standard Visium data (**Fig. 5c**). Moreover, we also tried estimating the WNT-signaling pathway activity by calculating a module score using a larger gene set of 42 genes ^25^, which detected enrichment of the pathway in the RRST data but not in the standard Visium data (**Fig. 5d**). These results highlight the importance of high quality data for molecular characterization of clinical samples, which for this particular sample could be achieved by RRST.

### RRST sheds light on cartilage and bone biology

Analysis of RNA profiles of cartilage and bone is a challenging task because cells in these tissues are embedded in dense extracellular matrices, which are also often mineralised ^26^. Extensive enzymatic digestion is typically required to isolate cells from these tissues, but the influence of such procedure on the transcriptional profiles of these cells is not fully understood, and whether sub-populations of cells remain in the undigested tissue is typically not reported ^27^. SRT offers a major advantage to study these tissues since gene expression can be analyzed without the need to isolate cells, together with the benefit of added spatial information.

The long-bones elongate via a process called endochondral ossification, in which streams of chondrocytes from the epiphyseal cartilage undergo successive differentiation stages and produce a mineralised cartilage matrix, which is subsequently remodeled and used as a scaffold on which new bone tissue is deposited ^28^. One of the later developmental stages in this process is the formation of a bony structure called the secondary ossification center (SOC) within the epiphyseal cartilage ^28^. In the proximal tibia of humans, this event occurs around birth ^29^, whereas in mice it is precisely determined to occur between postnatal day 7 and 11 ^30^. Within these few days, the SOC contains many different cell-types, including osteoblasts, hematopoietic cells, mesenchymal stromal cells and endothelial cells, which are suddenly located within a few cell-diameters of the resting-zone chondrocytes, potentially influencing these cells ^31^. To investigate potential effectors that derive from the newly forming SOC, we applied RRST to mouse growth plate specimens before SOC formation (postnatal day 4, P4) and immediately after SOC formation (postnatal day 11, P11) ^10^.

First, we aimed to benchmark our RRST protocol with the standard Visium protocol. In line with previous results on other tissue types processed in this study, we observed a 3-to 9-fold increase in the number of unique genes detected with RRST (**Fig. 6a**). Importantly, this trend was particularly clear in the cartilage and bone tissue, where we observed between 1298 to 1750 unique genes and between 2822 to 4000 UMIs on average with RRST, whereas the standard Visium protocol recovered less than 100 unique genes and UMIs on average (**Supplementary Fig. 11a,b,e and f**). The difference in the number of genes and UMIs was also evident in the surrounding tissues, and in addition we observed more even distribution of unique genes and UMIs in the RRST datasets (**Supplementary Fig. 11e** and **f**). Based on these observations, we decided to proceed with the higher quality RRST data for downstream analysis of the cartilage and bone tissue.

**Fig.6:**
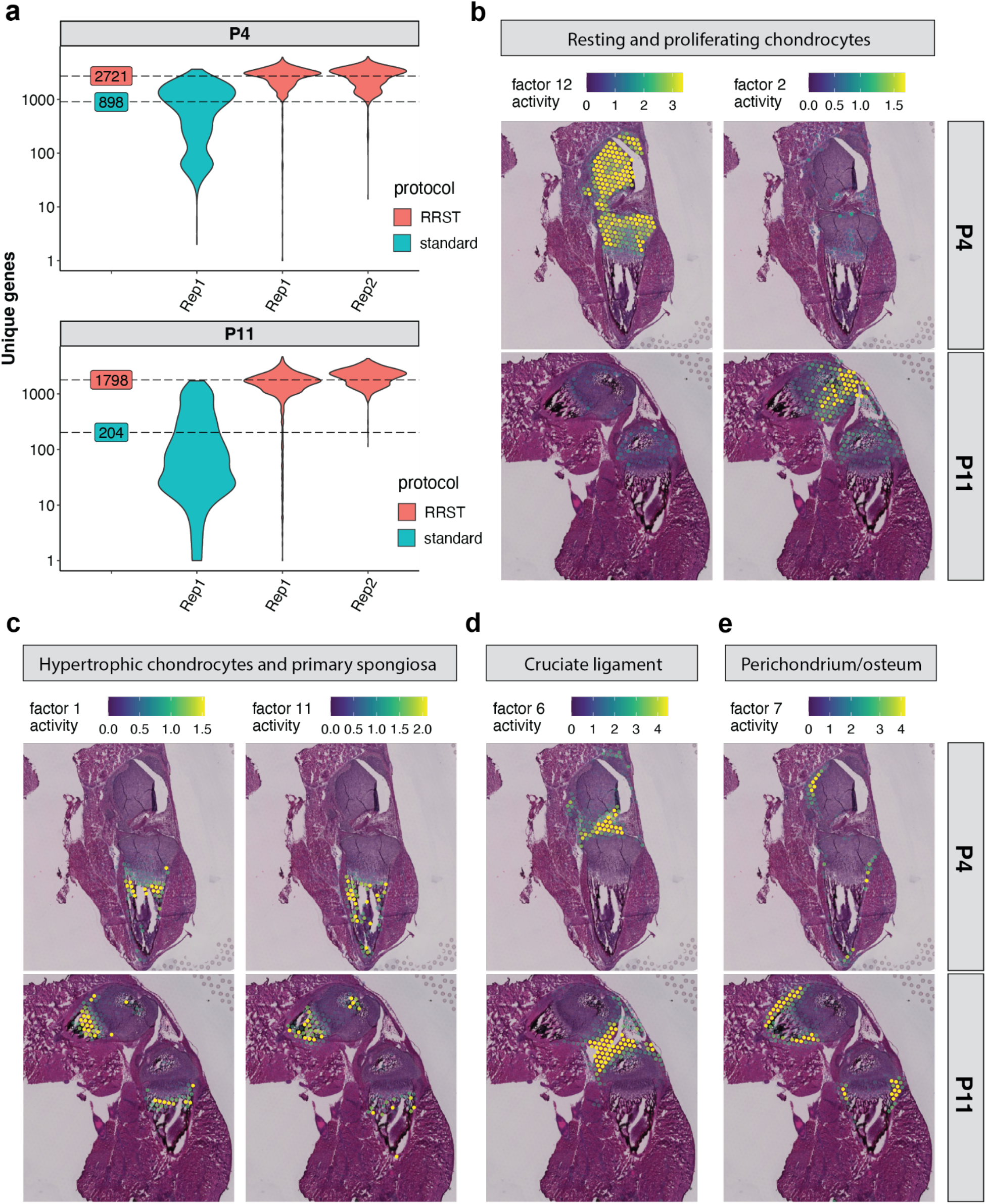
Comparison of standard Visium with RRST on mouse cartilage tissue. **a)** Average numbers of unique genes are highlighted by dashed lines for each protocol next to the violin plots. The y-axis represents log10-scaled counts. **b)** Following NNMF, Factors 12 and 2 associated with resting and proliferating chondrocytes. **c)** Factors 1 and 11 associated with hypertrophic chondrocytes and primary spongiosa. **d)** Factor 6 associated with the cruciate ligament. **e)** Factor 7 associated with Perichondrium and periosteum. Spot colors represent the factor activity, i.e. the contribution of each spot to the factor. The spot opacity has been scaled by the factor activity scores, making spots with lower scores more transparent.

Non-negative matrix factorization (NNMF) analysis identified several factors containing chondrocytes in the resting and proliferating zones (eg. Col2a1, Col9a1, **Fig. 6b** and **Supplementary Fig. 12a**) ^32^, hypertrophic chondrocytes and bone cells within the primary spongiosa (eg. Col10a1, Mmp9, Phospho1, Dmp1, Acp5, **Fig. 6c** and **Supplementary Fig. 12b**) ^33,34^, as well as the cruciate ligament (eg. Scx, Dkk3, **Fig. 6d** and **Supplementary Fig. 12c**) ^35,36^ and cells at the perichondrium/periosteum (eg. Thbs2, Tnn, **Fig. 6e** and **Supplementary Fig. 12d**) ^37^, which appeared in the distinct, expected anatomical locations. To explore possible secreted factors deriving from the newly forming SOC, we used the histological images to manually assign the spots within the cartilage into seven sub-clusters: “resting zone”, “proliferating zone”, “pre-hypertrophic”, “hypertrophic zone”, “SOC”, “SOC-adjacent resting zone” and those surrounding the cartilage that we grouped as “peripheral cells’’ (**Fig. 7a** and **Supplementary Fig. 11d**). To identify novel markers for these sub-clusters, we conducted differential gene expression analysis (**Fig. 7b**). We identified several genes specifically upregulated in the SOC and SOC adjacent zone; interestingly, one of these factors, Plxnd1, has previously been found to be expressed in newly forming ossification centers ^38^. Furthermore, we identified several soluble factors that were significantly upregulated within the SOC (namely Ccl9, Basp1 and Apln) and SOC-adjacent zone (Msmp). Thus, these results show that with RRST approach we open up an exciting possibility to gain deeper understanding of bone formation and other processes occuring in the skeleton in spatial context.

**Fig.7:**
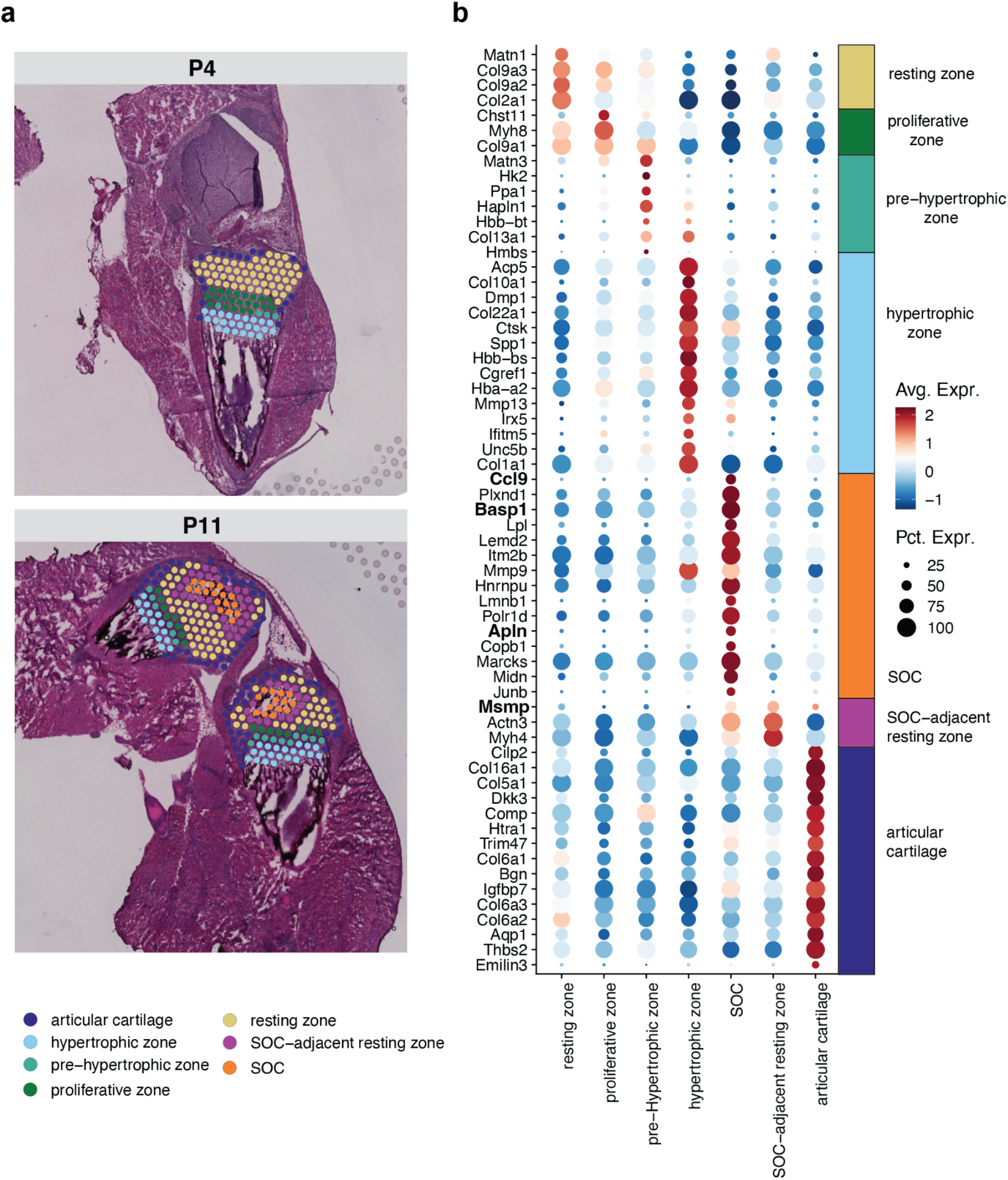
Investigating the potential secreted markers within mouse cartilage at 2 postnatal time points. **a)** Comparing the manually assigned sub-clusters between postnatal day 4 and day 11. **b)** Differentially up-regulated genes listed from highest to lowest fold change within each annotated region (avg_log2FC > 0.6 and a maximum of 15 genes per region).

## Discussion

Here we present the RNA Rescue Spatial Transcriptomics (RRST) profiling method, designed specifically for genome-wide spatial gene expression analysis of moderate to low quality fresh frozen (FF) samples. Recent developments in the field have made it possible to generate SRT data from FFPE samples, which is the preferred fixation method for storing biological material in biobanks. Formalin-fixation provides better preservation of morphology and makes the material compatible with spatial mRNA-protein co-detection assays. While FFPE sample preservation has its advantages, overfixation leading to heavily crosslinked RNA is a common issue, which may introduce biases in the analysis of both RNA and DNA in those samples ^39^. Hence, we modified the commercially available Visium FFPE spatial gene expression protocol to be applicable on FF tissues by introducing three modifications: (1) a short formalin fixation step to make RRST compatible with Visium FFPE protocol, (2) a baking step for reinforced tissue section adhesion and prevention of detachment and (3) removal of the crosslink-reversal step, which shortens the overall protocol time. In addition, we believe that RRST will increase flexibility for researchers working with snap-frozen samples, in particular to make SRT compatible with other modalities that rely on FF specimens such as single nuclei sequencing or mass spectrometry in order to obtain paired data from the same tissue block.

In this work, we analyzed 52 tissue sections across 7 different tissue types to demonstrate the versatility of RRST protocol. Although standard Visium protocol, which relies on methanol-fixation, has been shown to work in high quality FF specimens, our analysis of mouse brain and prostate cancer tissue demonstrates that RRST performs equally well in tissues with high RIN values and exhibits better performance in low-quality samples as demonstrated by the increased number of detected genes and transcripts in several different tissue types. We show that in samples collected from the human small intestine and colon, we observed severe RNA degradation in epithelial tissues; however, with the RRST protocol we were able to recover spatial data from these tissues when the standard protocol failed.

Notably, RRST allowed us to identify characteristic WNT-signaling pathway genes in a medulloblastoma WNT subtype of pediatric brain tumors, which would have been otherwise overlooked in standard Visium-derived data. Moreover, the RRST protocol does not require tissue optimization, making it advantageous in situations where little material is available as is often the case with precious clinical specimens. In addition, we demonstrate that RRST protocol can successfully generate transcriptomic profiles in challenging tissue types such as adult human lung or mouse cartilage/bone. For example, by applying RRST to adult human lung tissue we are able to provide a more detailed, data-driven characterization of different tissue compartments. The additional information that we observe in the RRST data makes the technology more relevant for studies of the respiratory system.

To the best of our knowledge, we have generated the first spatially resolved transcriptomics dataset from cartilage and bone tissue, which opens up new possibilities to study the composition and communication of cells in the skeletal system for example to better understand cellular microenvironments within the bone marrow ^40^, the crucial gradient of cell identities at attachment sites between muscle and bone ^34^, as well as to study diseases such as osteoarthritis whose step-wise progressive degeneration involves complex interplay between various tissues, including cartilage and bone ^41,42^. We demonstrate that by applying RRST to mouse cartilage/bone tissue, we could identify four soluble factors expressed within the SOC or the SOC-adjacent zone, which have the potential to influence the chondrocytes, based on their close proximity. Since Apln has been shown to be involved in endothelial cell activation during angiogenesis ^43^, and Ccl9 in the maturation of osteoclasts ^44^, their expression may reflect the ongoing growth and remodeling of the SOC. However, further research is required to reveal the precise roles of the four soluble factors in the SOC and their possible influence on bone growth.

In summary, we show that our RRST protocol recovers higher amounts of mRNA than the standard Visium protocol from degraded or otherwise challenging FF tissue blocks. Taken together, our results indicate that RRST is a powerful and versatile method, which can be used to accelerate discoveries in developmental biology, disease pathology, and clinical translational research.

## Methods

### Ethics declaration

The study was performed according to the Declaration of Helsinki, Basel Declaration and Good Clinical Practice. All human subjects were provided with full and adequate verbal and written information about the study before their participation. Written informed consent was obtained from all participating subjects before enrolment in the study.

Use of prostate cancer samples was approved by the Regional Ethical Review Board (REPN) Uppsala, Sweden before study initiation (Dnr 2011/066/2, Landstinget Västmanland, Sari Stenius). Lung samples were obtained from deceased donors by the Cambridge Biorepository for Translational Medicine (CBTM) with informed consent from the donor families and approval from the NRES Committee of East of England – Cambridge South (15/EE/0152), the project has received funding from the European Union’s Horizon 2020 research and innovation programme under a grant agreement (no. 874656, discovAIR).

GI tract specimens were approved by the medical ethics committee of University Hospitals Leuven (approval no. S62935).

Use of pediatric brain tumor samples was approved by the Regional Ethical Review Board (EPN), Stockholm, Sweden (DNR 2018/3-31, Monica Nister).

Mouse bone samples were collected according to DNR 16673/2020, approved by Stockholm’s animal experiment ethics committee (Stockholms djurförsöksetiska nämnd).

Mouse brain sample was purchased from Adlego Biomedical company, that operates under ethical permission nr. 17114-2020.

### Samples information

#### Mouse brain

A mouse brain sample was selected from a batch of commercially purchased specimens from Adlego Biomedical.

#### Prostate cancer sample

Prostate cancer sample was obtained from a surgically removed prostate at Västerås Hospital in Sweden.

#### Lung specimens

Postmortem samples from lung tissue were collected at the department of Molecular Biosciences, Science for Life Laboratory, Stockholm, Sweden. Autopsy samples were selected from two healthy donors.

#### GI specimens

Samples were collected from patients undergoing colorectal surgery. Collection of small intestine and colon samples biopsies was performed at the department of Chronic Disease and Metabolism, Katholieke Universiteit Leuven, Belgium.

#### Pediatric brain tumor samples

Samples were obtained from The Swedish Childhood Tumor Biobank.

#### Mouse cartilage/bone

Tissues were collected from postnatal mice at four and eleven days of age. Briefly, hind-limbs were dissected, the skin and surrounding soft tissues were quickly trimmed. Femora and tibiae were dissected through the diaphysis and the tissue including the knee joint, proximal tibia and distal femur (with remaining soft tissues) was embedded into OCT in a cryomold. The samples were rapidly frozen using a hexane bath.

### RNA quality evaluation

RNA was extracted from tissue sections using the RNeasy Mini kit (Qiagen). RINs were measured using the Agilent Bioanalyzer.

### Standard Visium Spatial Gene Expression library preparation

Fresh-frozen samples were cryo-sectioned at 10 μm thickness, placed onto Visium glass slides and stored in −80°C before processing. Spatial gene expression libraries were generated following 10X Genomic Visium Spatial Gene Expression protocol (User Guide, CG000239 Rev F). Libraries were sequenced on Nextseq2000 (Illumina). Length of read 1 was 28 bp and read 2 150 bp.

### RRST Gene Expression library preparation

The fresh-frozen samples were cryo-sectioned at 10 μm thickness, placed onto Visium glass slides and stored in −80°C before processing. Visium slides were taken out of the −80°C freezer and placed on a thermocycler pre-heated at 37°C for 1 minute, followed by immediate fixation in 4% methanol-free formaldehyde (thermofisher, Catalog number: 28906) solution for 10 minutes at room temperature. After fixation, Visium slides were washed twice in 1xPBS, then heated up to 37°C for 20 minutes using a thermocycler, cooled down to room temperature and followed by Hematoxylin-Eosin staining optimized for each tissue type and imaging. Directly after imaging, slides were washed with MQ water, air-dried and placed inside the plastic Visium cassette. Sections were treated with 0.1N HCl for 1 minute at room temperature, and washed in 1xPBS. The Visium Spatial Gene Expression for FFPE reagent kit (10x Genomics, Pleasanton, CA, USA) was used for the downstream steps. Decrosslinking step was skipped, immediately proceeding with probe Pre-hybridization step for 15 minutes at room temperature, followed by Probe Hybridization overnight according to 10X Visium Spatial Gene Expression Reagent Kits for FFPE protocol and the rest of the library preparation (User Guide, CG000407 Rev C). Finished libraries were sequenced on Nextseq2000 (Illumina). Length of read 1 and read 2 were 28 base pairs and 50 base pairs, respectively.

### Data processing

Sequenced libraries were processed using Space Ranger software (version 1.2.1 for standard Visium data and version 1.3.1 for RRST data, 10X Genomics). Reads were aligned to the pre-built human or mouse reference genome provided by 10x Genomics (GRCh38 for human data or mm10 for mouse data, version 32, ensembl 98), which includes a GTF file, a fasta file and a STAR index.

### Data filtering and pre-processing

Processing and analysis of spatial transcriptomics data obtained with either RRST or standard Visium was performed using R (v4.1.3) and the single-cell genomics toolkit *Seurat* and the spatial transcriptomics toolkit *STUtility*. Adult human colon and small intestine data was manually annotated into major tissue compartments based on tissue morphology (H&E image) using the interactive shiny app provided with the ManualAnnotation function in *STUtility*. Adult human colon data was categorized into three groups: “mucosa”, “submucosa” and “muscularis” whereas small intestine data was categorized into 5 groups: “mucosa”, “TLS”, “submucosa”, “muscularis” and “serosa”. **Table 1** provides a summary of the filtering settings used for each dataset. Detailed instructions for each sample type are provided in the section below.

**Table 1:** Overview of spatially resolved transcriptomics samples and filtering settings used in preprocessing steps.

| dataset | # RRST samples | # standard Visium samples | # biological replicates | filter |
| --- | --- | --- | --- | --- |
| mouse brain | 4 | 4 | 1 | no filter |
| human prostate cancer | 2 | 2 | 1 | no filter |
| adult human lung | 2 | 4 | 2 | keep spots with > 300 unique genes |
| adult human colon | 4 | 2 | 2 | keep spots annotated as “mucosa”, “submucosa” or “muscularis” |
| adult human small intestine | 2 | 12 | 1 | keep spots annotated as “mucosa”, “TLS”, “submucosa”, “muscularis” or “serosa” and spots with > 100 unique genes |
| mouse bone | 2 | 4 | 2 | keep spots with > 500 unique genes |
| pediatric brain tumor | 4 | 4 | 2 | no filter |

#### Mouse brain and human prostate cancer

A total of 8 mouse brain tissue sections (4xRRST and 4xstandard) and 4 prostate cancer tissue sections (2xRRST and 2xstandard) were used for the analysis. Spatial visualization of unique genes were created using the ST.FeaturePlot function (*STUtility*) and violin plots with the *ggplot2* R package. The median number of unique genes were calculated for each protocol and sample and visualized next to the violin plots. Gene-gene scatter plots comparing log-transformed UMI counts were created as follows: (1) raw expression matrices were extracted for each data type (RRST or standard Visium) followed by aggregating the expression values for each gene, (2) aggregated expression values were log-transformed with a pseudocount of 1 (log1p). Pearson R scores and p-values were calculated using the stat_cor function from the *ggpubr* R package. Gene-gene scatter plots comparing detection rates were created as follows: (1) raw expression matrices were extracted for each data type (RRST or standard Visium) and the detection rates were estimated for each gene as the proportion of spots with detected UMI counts.

#### Adult human lung

A total of 6 adult human lung tissue sections (2xRRST and 4xstandard), collected from two samples were used for the analysis. After filtering out spots with fewer than 301 unique genes detected, the data was normalized and subjected to a basic analysis workflow using functions from the *Seurat* R package. The filtered data was split by sample (LNG1 and LNG1), which were analyzed separately. Normalization and scaling of the data was conducted using the NormalizeData and ScaleData functions. The top 2000 most variable genes were detected using the vst method (FindVariableFeatures) followed by dimensionality reduction by PCA (RunPCA). A shared nearest neighbor (SNN) graph was constructed based on the first 20 principal components (FindNeighbors) followed by graph-based clustering with the resolution parameter set to 0.8 (FindClusters). Finally, a Uniform Manifold Approximation and Projection (UMAP) embedding was computed based on the first 20 principal components (RunUMAP, min.dist = 0.3, n.epochs = 1000). Marker detection was conducted by calculating differential expression for each cluster against the background (remaining clusters) with a log fold change threshold of 0.25 and an adjusted p-value threshold of 0.01 using the FindAllMarkers function. Cluster annotations were assigned based on the expression of canonical markers (obtained from a scRNA-seq atlas of the human lung ^45^) and spatial co-localization with histological landmarks.

#### Adult human colon

A total of 6 adult human colon Visium datasets (4xRRST and 2xstandard), obtained from 2 samples were used for the analysis. Spots in these datasets were manually labeled using the ManualAnnotation function from *STUtility* into three major regions based on histology: “mucosa”, “submucosa” and “muscularis”. Unlabeled spots were removed prior to downstream analysis using the SubsetSTData function from *STUtility*. Datasets 2, 3, 4 and 6 were used for the spatial plots in **Fig. 3a, b** (see **Table 1**). Violin plots showing the distribution of unique genes in the three major regions were created for all 6 datasets using the *ggplot2* R package. Prior to normalization, the data was filtered to only keep genes expressed in both RRST and standard data. The dataset was then normalized using the NormalizeData function from *Seurat*. 11 intestinal epithelial marker genes were selected based on two criteria: (1) high spatial variability in the spatially resolved transcriptomics data (the data presented here), and (2) high differential expression in epithelial cells identified in the Gut Cell Atlas ^19^. The normalized expression of these 11 intestinal epithelial marker genes were then visualized as violin plots for spots annotated as “mucosa”.

#### Human small intestine

A total of 14 adult human small intestine Visium datasets (2xRRST and 12xstandard), obtained from a single specimen collected over a time span of ~2 years, were used for the analysis. Spots in these datasets were manually labeled using the ManualAnnotation function from *STUtility* into five major regions based on histology: “mucosa”, “TLS”, “submucosa”, “muscularis” and “serosa”. Unlabeled spots were removed prior to downstream analysis using the SubsetSTData function from *STUtility*. Violin plots showing the distribution of unique genes in the five major regions were created for all 14 datasets using the *ggplot2* R package, with the average number of unique genes highlighted for each time point. The biotype content was calculated for 10 biotypes: IG(C|J|V), TR(C|J|V), lincRNA, protein coding, mitochondrial protein coding and ribosomal protein coding genes. All other transcripts were labeled as “other”. For each biotype and within each time point, a percentage was calculated by dividing the UMIs for the biotype with the total number of UMIs. The gene annotations were obtained from the GTF file used for mapping with spaceranger. Note that the RRST protocol only targets protein coding transcripts, immunoglobulin transcripts and T-cell receptor transcripts. Next, we split the dataset by time points, filtered out spots with less than or equal to 100 unique genes, and normalized each subset with the NormalizeData function. For differential expression analysis of the mucosa, we used the FindMarkers function to identify marker genes with a log fold change threshold of 0.25 and an adjusted p-value lower than 0.01 (max.cells.per.ident = 1000, ident.1 = “mucosa”, only.pos = TRUE). After filtering the differentially expressed genes (DEGs) based on log fold change and adjusted p-values, the results were summarized in an “upset” plot highlighting genes that were differentially expressed at single or multiple time points. 6 intestinal epithelial marker genes were selected based on two criteria: (1) high spatial variability in the spatially resolved transcriptomics data (the data presented here), and (2) high differential expression in epithelial cells identified in the Gut Cell Atlas ^19^. The normalized expression of these 6 intestinal epithelial marker genes were then visualized as spatial maps with ST.FeaturePlot (*STUtility*) in 3 selected tissue sections, 1 from each time point.

#### Pediatric brain tumor

A total of 8 pediatric brain tumor tissue sections (4xRRST and 4xstandard), collected from two tissue blocks (medulloblastoma and NOS subtypes), were used for the analysis. The distribution of unique genes for all 8 tissue sections were visualized as violin plots colored by protocol, and with the average number of unique genes highlighted next to the violin plots. 1 representative tissue section was selected from each combination of protocol and sample to show the distribution of unique genes together with the corresponding H&E image. Next, we downloaded cancer hallmark gene sets from MsigDB ^46,47^ for WNT β-catenin-signaling and TGFβ-signaling. These gene sets were then used to compute enrichment scores from the normalized medulloblastoma data with the AddModuleScore function from *Seurat*. These module scores were then visualized as spatial maps on 1 representative tissue section from each protocol (RRST or standard). Next, we selected 6 known WNT-signaling marker genes and visualized their normalized expression distributions as violin plots in the medulloblastoma data.

#### Mouse bone

A total of 6 tissue sections (4xRRST and 2xstandard), collected from two tissue blocks (P4 and P11), were used for the comparison shown in **Figure 6a** and **Supplementary Fig. 11**. The spots were manually annotated into two regions: “cartilage/bone” and “surrounding” tissue. Distributions of unique genes and UMIs at the two post-natal stages and in manually annotated regions (split by protocol) were visualized with violin plots using the *ggplot2* R package and spatial maps were created with the FeatureOverlay function from *STUtility*. Only the RRST samples were used for subsequent data analysis. First, the “cartilage/bone” region was manually annotated into seven sub regions: “resting zone”, “proliferative zone”, “pre-Hypertrophic zone”, “hypertrophic zone”, “SOC”, “SOC-adjacent resting zone” and “articular cartilage” (shown in **Fig. 7a**). Spots with at least 500 unique genes were kept prior to normalization using variance stabilizing transformation (vst) implemented in the SCTransform function from *Seurat*. The Non-Negative Matrix Factorization (NNMF) was computed on the filtered and normalized data using the RunNMF function from *STUtility*, with the number of factors set to 30. Based on visual inspection we identified 8 factors colocalized with various structures of the cartilage/bone tissue region: factor_12, factor_2, factor_1, factor_11, factor_6 and factor_7 (shown in **Fig. 6b, c, d, e** and **Supplementary Fig. 12**). Next, we created a subset of the data including only the seven sub regions defined within the cartilage/bone, with the goal of extracting marker genes from each sub region by differential expression analysis (DEA). Prior to running the DEA, we first renormalized the raw UMI counts with the NormalizeData and ScaleData functions. The DEA was conducted using FindAllMarkers from *Seurat*, while filtering out genes with adjusted p-values lower than 0.01 and average log fold change values higher than 0.25. Marker genes visualized in **Fig. 7** were selected by keeping those with average log fold change values higher than 0.6 and maximum 15 genes per sub region.

## Data and code availability

All data required to replicate the analyses, including spaceranger output files, H&E images and additional files will be available at Mendeley Data. Sequence data for all samples presented in this study will be available upon publication.

## Acknowledgment

This project has received funding from the European Research Council (ERC) under the European Union’s Horizon 2020 research and innovation programme (Discovair, grant agreement No. 782 101021019). The study was also supported by The Swedish Cancer Society, Swedish Foundation for Strategic Research, The Leona M. and Harry B. Helmsley Charitable Trust, Swedish Childhood Cancer Fund and Science for Life Laboratory. We would like to thank Krishnaa Mahbubani for collecting human lung samples and the National Genomics Infrastructure (NGI), Sweden for providing infrastructure support. We thank Drs. Annelie Mollbrink, Alma Andersson and Marco Vicari for helpful assistance, discussions and reading the manuscript.

## Author contributions

R.M, Z.A., L.L., initiated the project; R.M, Z.A., L.A.G. and X.M.A. planned and performed the experiments; L.L. analyzed all the data and generated the figures. L.K. helped analyze the pediatric brain tumor samples; P.T.N and M.A. provided mouse bone samples, led the bone/cartilage biology part and wrote the relevant sections; A.S. undertook histopathological analysis; G.B., A.D.S. and N.S. provided adult human colon and small intestine samples; N. Schultz. provided the prostate sample. M.N. provided pediatric brain tumor samples; C.S. and A.F. provided lung samples. A.J. provided advice; R.M, Z.A., L.L., drafted the manuscript; all authors read and approved the final manuscript. J.L. provided project guidance and supervision. L.A.G. and X.M.A. contributed equally to this work.

## Supplementary

**Supplementary Table 1:** Samples used in this study with their corresponding ID, RNA Integrity Number (RIN) and DV200.

| Sample Type | Sample ID | RIN | DV200 (%) |
| --- | --- | --- | --- |
| Mouse brain | MB | 8.8 | 93 |
| Prostate cancer | PC | 10 | 100 |
| Human lung | LNG1 | 7.1 | 90 |
| Human lung | LNG2 | 6.8 | 89 |
| Human colon | CLN1 | 5.1 | 77 |
| Human colon | CLN2 | 4.5 | 77 |
| Human small intestine | SI | 7.8 | 86 |
| Pediatric brain tumor | WNT medulloblastoma (PBT1) | 7.1 | 90 |
| Pediatric brain tumor | NOS (PBT2) | 7 | 88 |
| Mouse bone | P4 | 6.2 | 87 |
| Mouse bone | P11 | 5.6 | 82 |

**Supplementary Figure 1:**
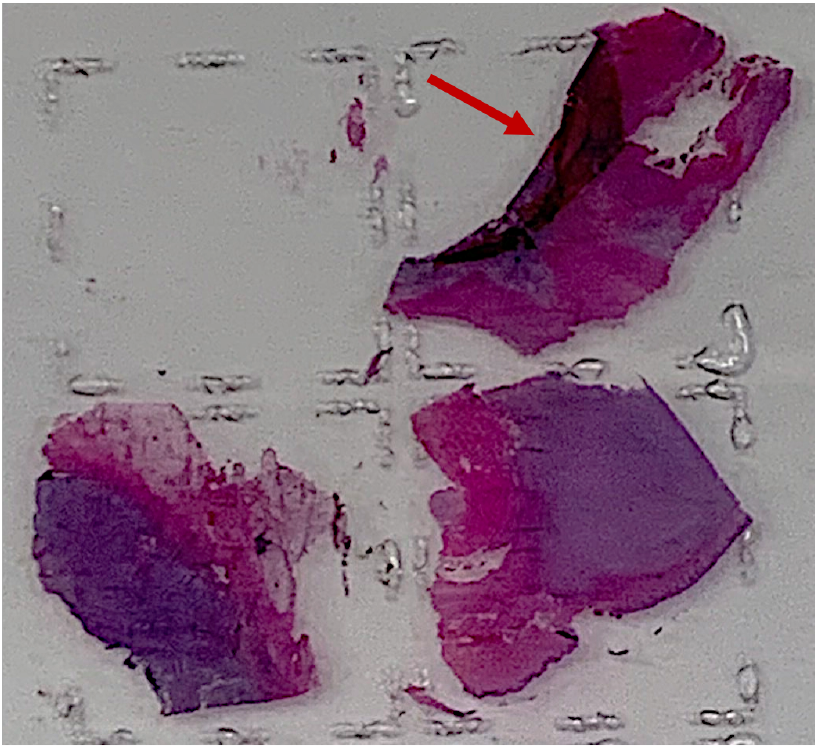
Tissue detachment. Tissue section detachment from a spatial gene expression array indicated by an arrow right after staining steps. **(plus Supplementary Video 1)**.

**Supplementary Figure 2:**
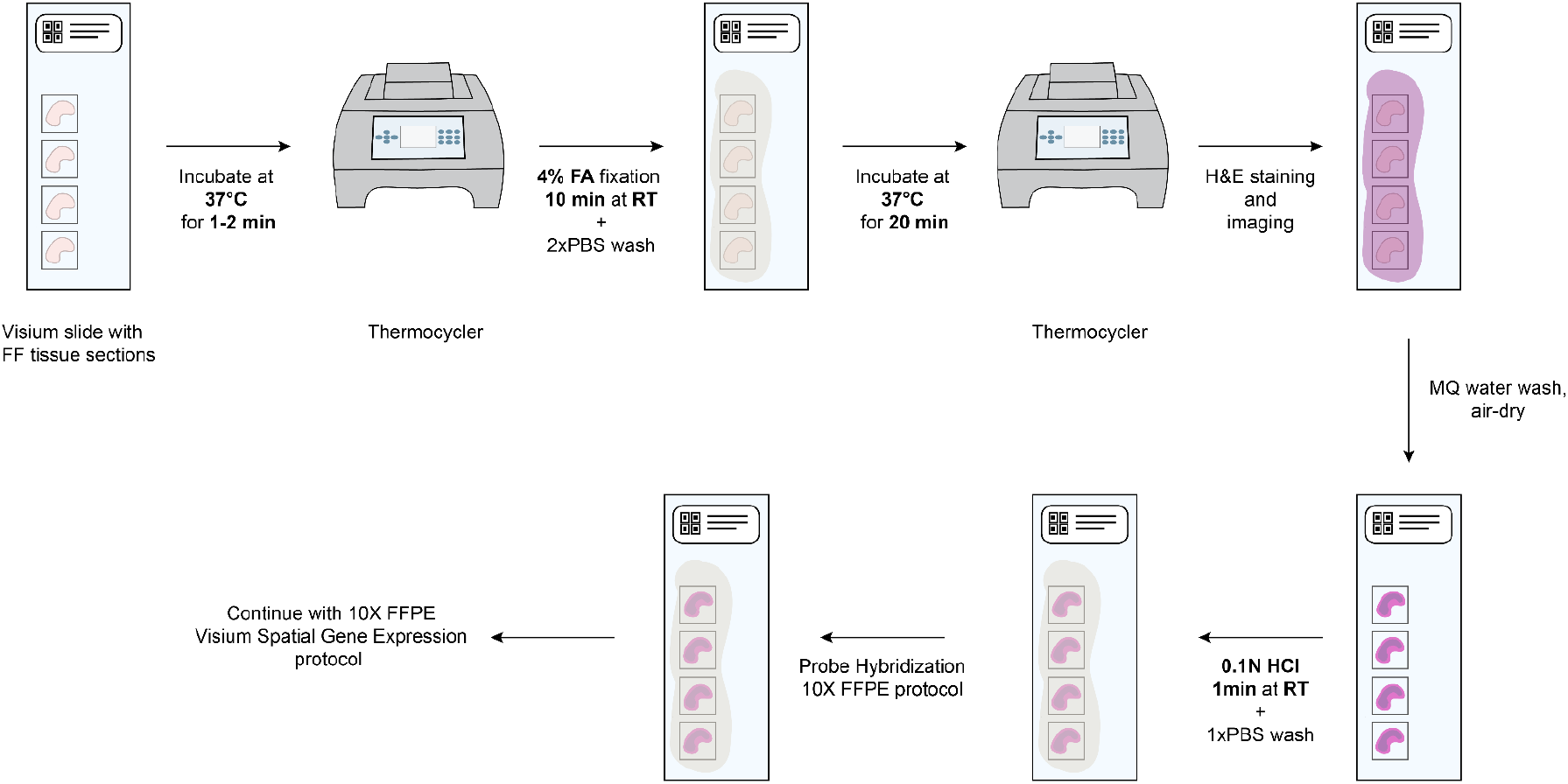
Schematic overview of the RRST protocol workflow.

**Supplementary Figure 3:**
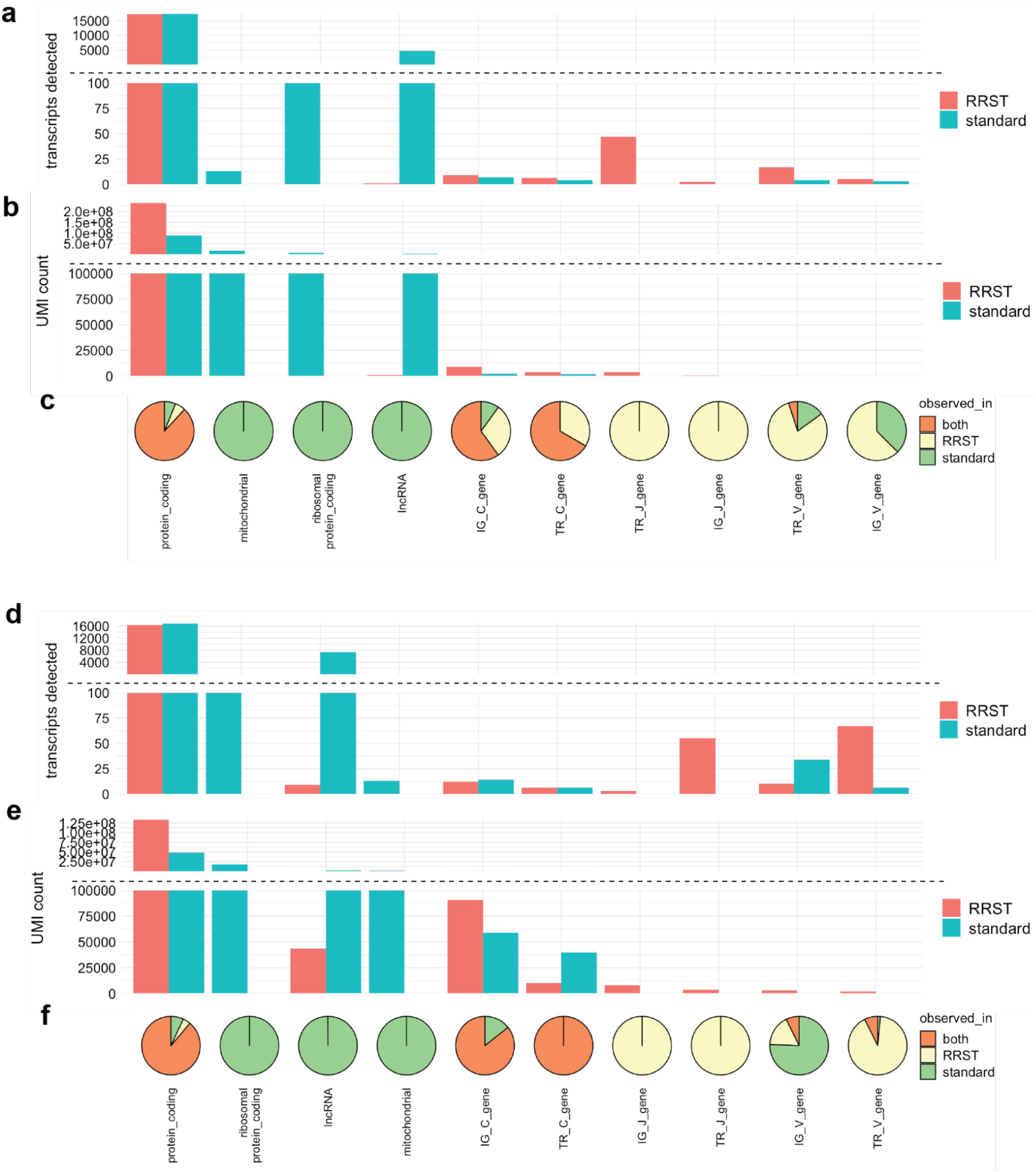
Biotype of transcripts present in Visium and RRST mouse brain and prostate cancer datasets. **a, b)** Transcripts detected (**a**) and UMI counts (**b**) for each biotype in the mouse brain data (RRST n=4, standard n=4). The fill color of the bars corresponds to the protocol used. The y-axis has been cut to display bio types with fewer detected transcripts or UMI counts. **c)** Pie charts below each bar in **b** display the proportions of transcripts that are detected with both methods, RRST only or standard Visium only. For example, lncRNA, mitochondrial transcripts and ribosomal protein coding gene transcripts are only detected with the standard Visium protocol. **d, e)** Transcripts detected (**d**) and UMI counts (**e**) for each biotype in the prostate cancer data (RRST n=2, standard n=2). The fill color of the bars corresponds to the protocol used. The y-axis has been cut to display bio types with fewer detected transcripts or UMI counts. **f)** Pie charts below each bar in **e** display the proportions of transcripts that are detected with both methods, RRST only or standard Visium only.

**Supplementary Figure 4:**
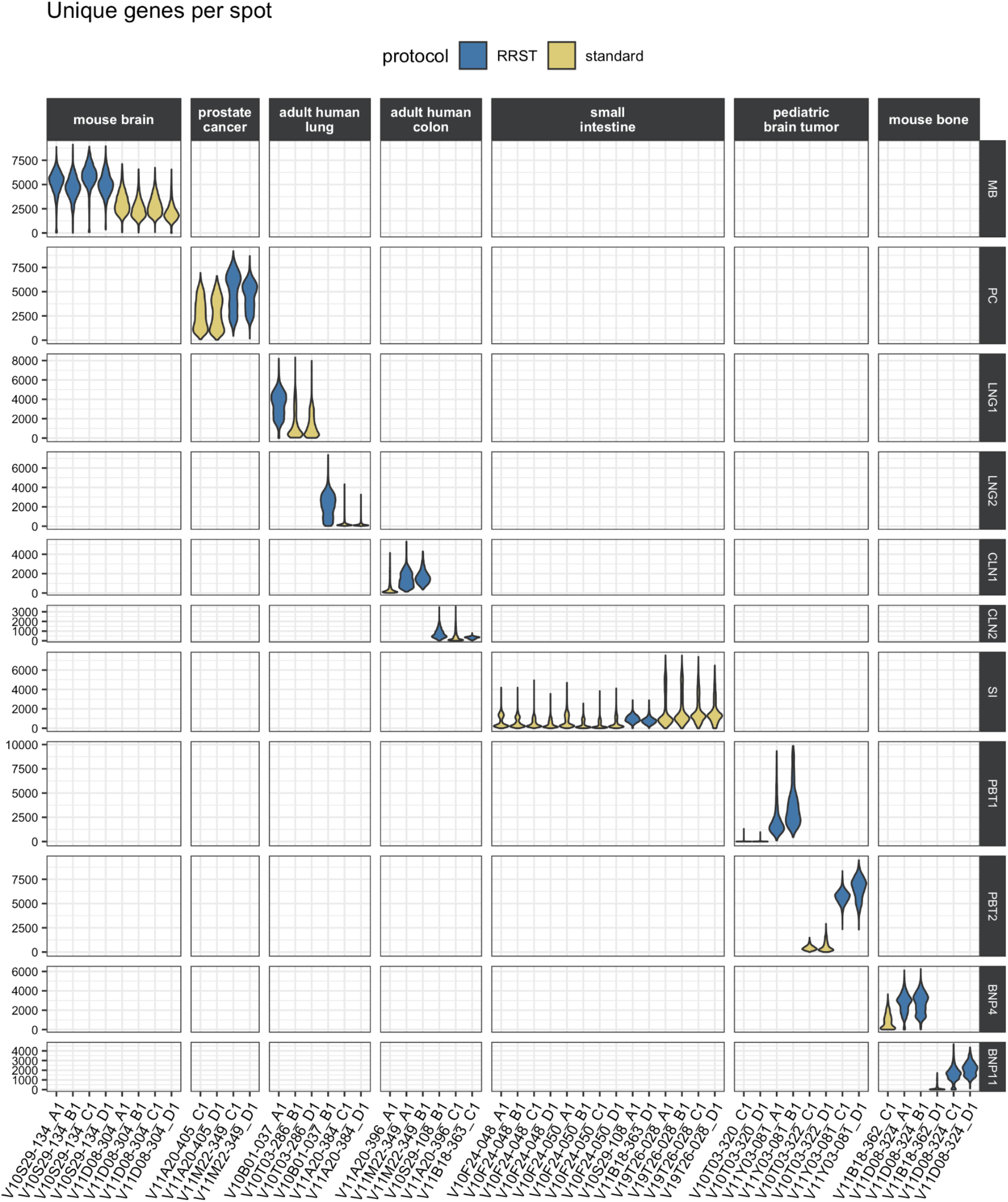
Distribution of unique genes per spot for all SRT datasets included in this study visualized as violin plots. The fill color of the violin plots indicates the protocol used to generate the data. Columns are sorted by sample origin and rows are sorted by sample ID. MB, mouse brain; PC, prostate cancer; LNG, adult human lung; CLN, adult human colon; SI, adult human small intestine; PBT, pediatric brain tumor; BN, mouse bone.

**Supplementary Fig.5:**
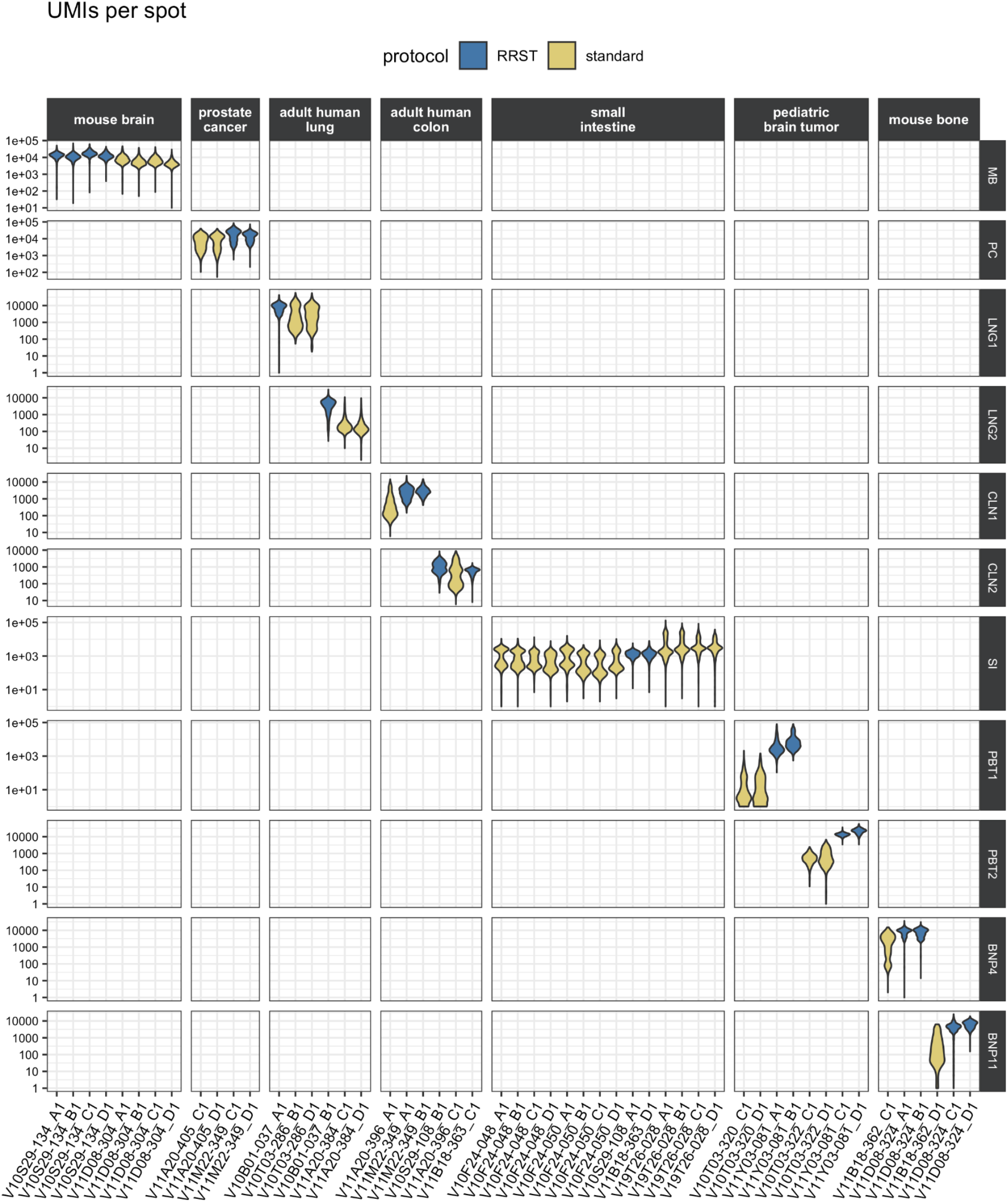
Distribution of UMIs per spot for all SRT datasets included in this study visualized as violin plots. The fill color of the violin plots indicates the protocol used to generate the data. Columns are sorted by sample origin and rows are sorted by sample ID. MB, mouse brain; PC, prostate cancer; LNG, adult human lung; CLN, adult human colon; SI, adult human small intestine; PBT, pediatric brain tumor; BN, mouse bone.

**Supplementary Figure 6:**
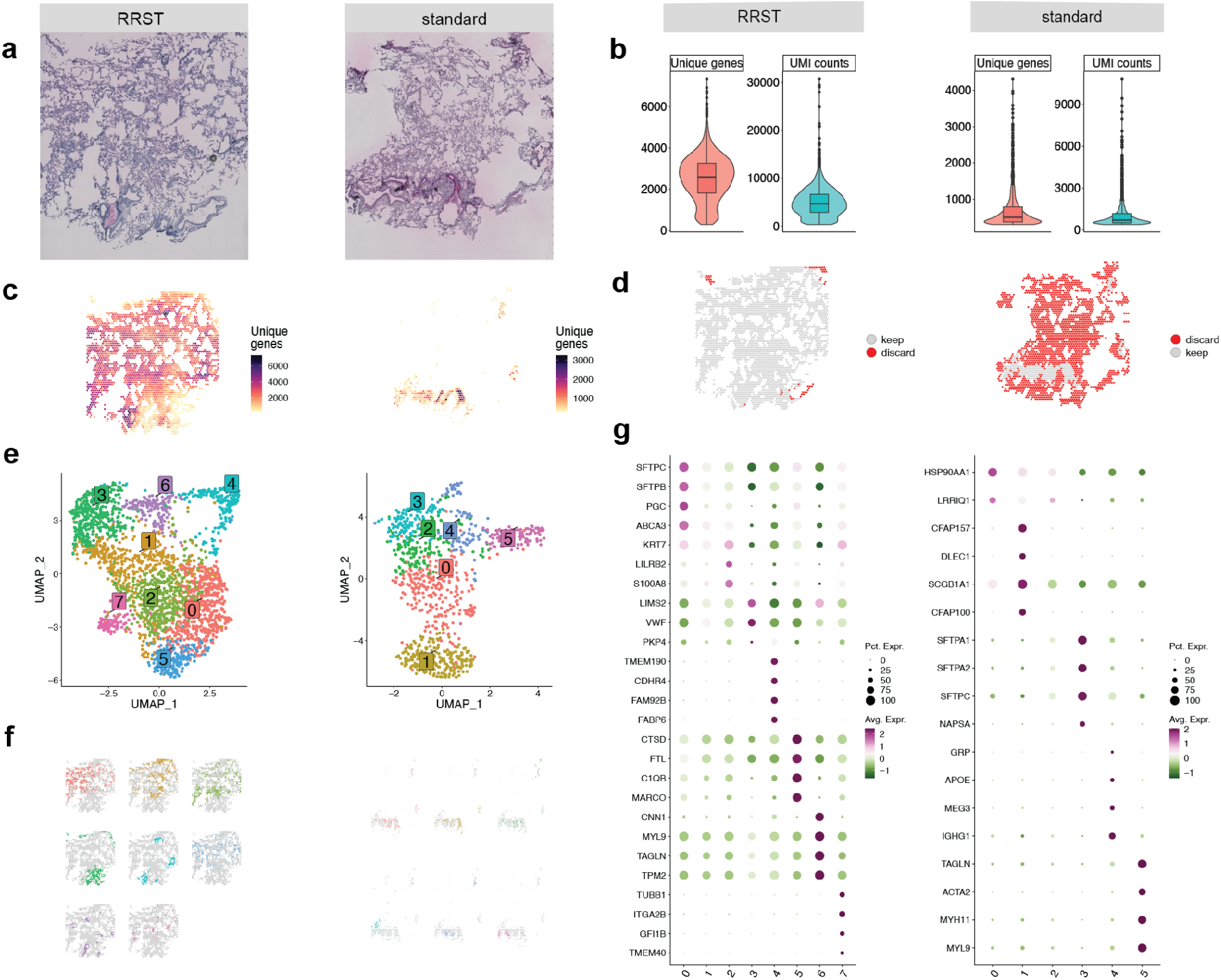
Comparison of RRST and standard Visium in human adult lung tissue (LNG2). Each subplot shows the RRST data on the left side and the standard Visium data on the right side. **a)** H&E images of two representative tissue sections from the same tissue block. **b)** Violin plots showing the distribution of unique genes and UMI counts for the adult lung data. **c)** Unique genes per spot mapped on tissue coordinates. **d)** Spatial visualization showing what spots were discarded due to low quality (less than 300 unique genes detected). **e)** UMAP embedding of adult lung data colored by clusters detected by unsupervised graph-based clustering (louvain). **f)** Split view of clusters (same as in **e**) mapped on tissue coordinates. **g)** Dot plots of the top marker genes for each cluster.

**Supplementary Figure 7:**
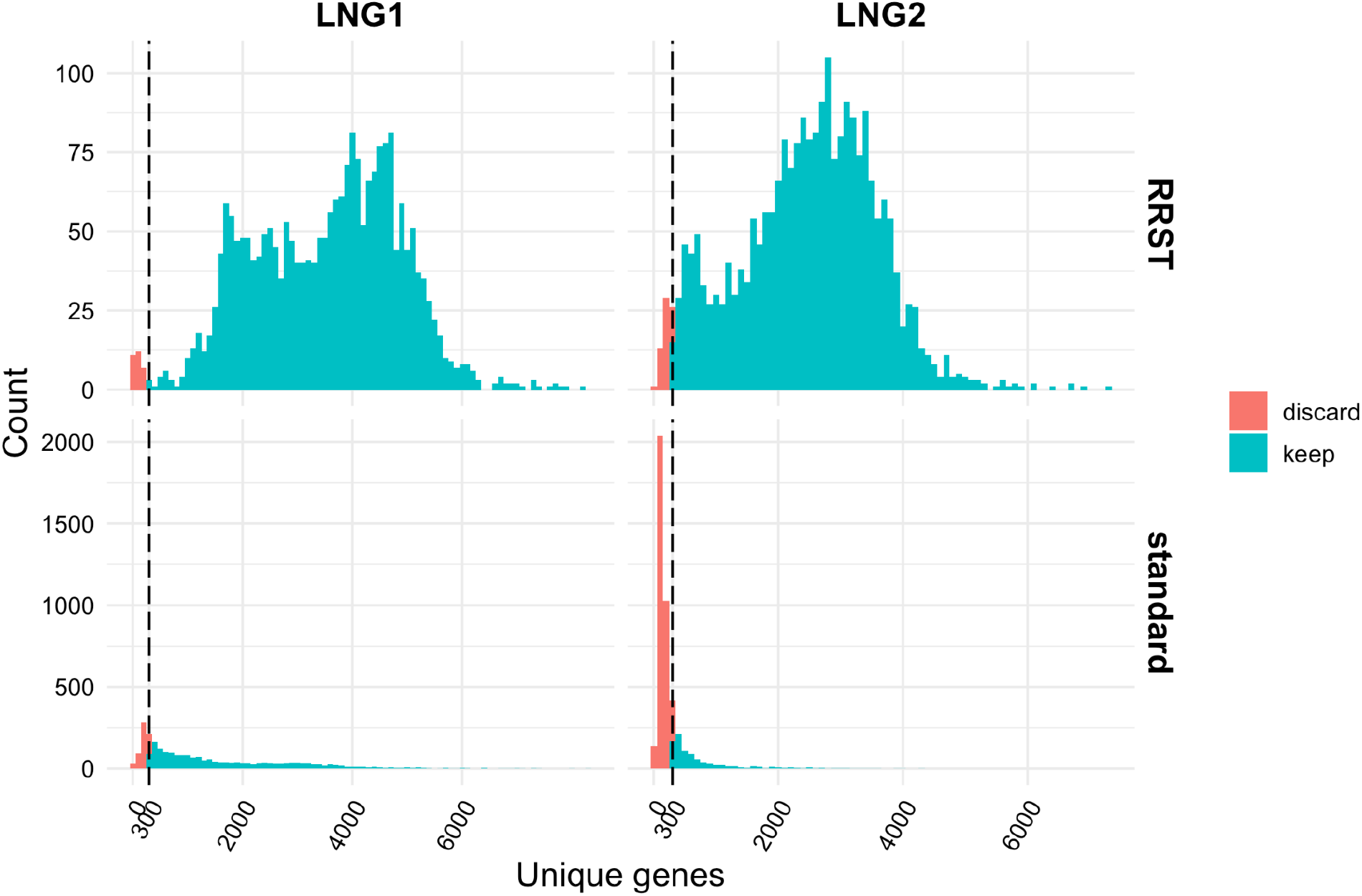
Histograms showing the distribution of unique genes per spot in lung tissue data obtained with RRST and standard Visium. A cutoff threshold of 300 was used as a filtering threshold to remove low quality spots. Blue bars represent spots that passed the cutoff threshold whereas red bars represent spots that were discarded. ~1.3% - 2.3% of spots were discarded from the two RRST datasets and ~21%-81% of spots were discarded for the standard Visium dataset.

**Supplementary Figure 8:**
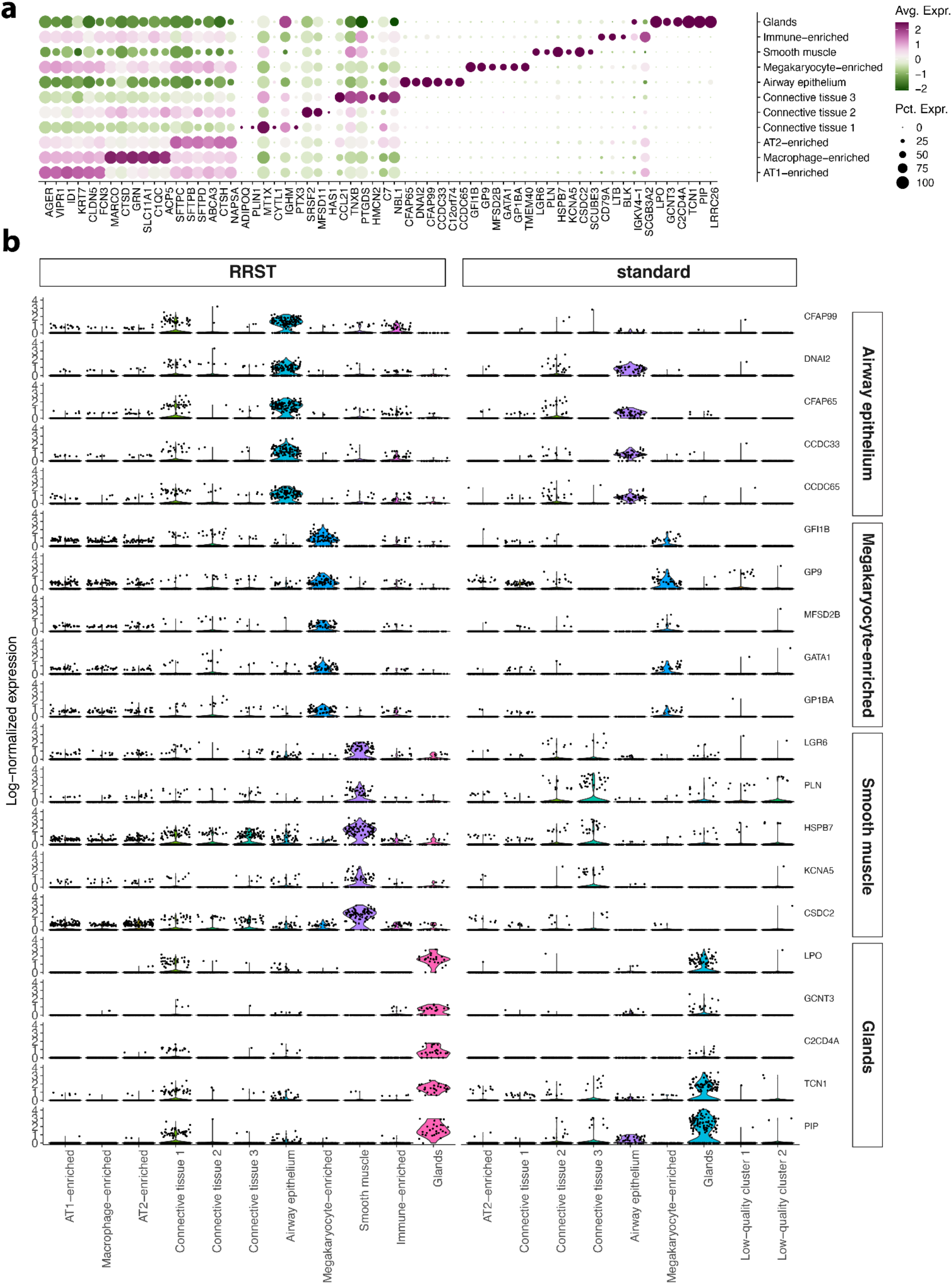
Visualization of marker genes detected in an adult human lung (LNG1) RRST dataset. **a)** Top marker genes for clusters detected in LNG1 RRST data. Dot colors correspond to averaged log-normalized and scaled gene expression (Avg. Expr.), and dot sizes correspond to the percentage of spots where the gene is detected (Pct. Expr.). **b)** Visualization of selected marker genes derived from data-driven clustering and differential expression analysis of LNG1 data. The left panel represents data obtained with RRST and the right panel represents data obtained with the standard Visium protocol. Five markers for each of four different tissue types are highlighted: airway epithelium, megakaryocyte-enriched, smooth muscle and glands.

**Supplementary Figure 9:**
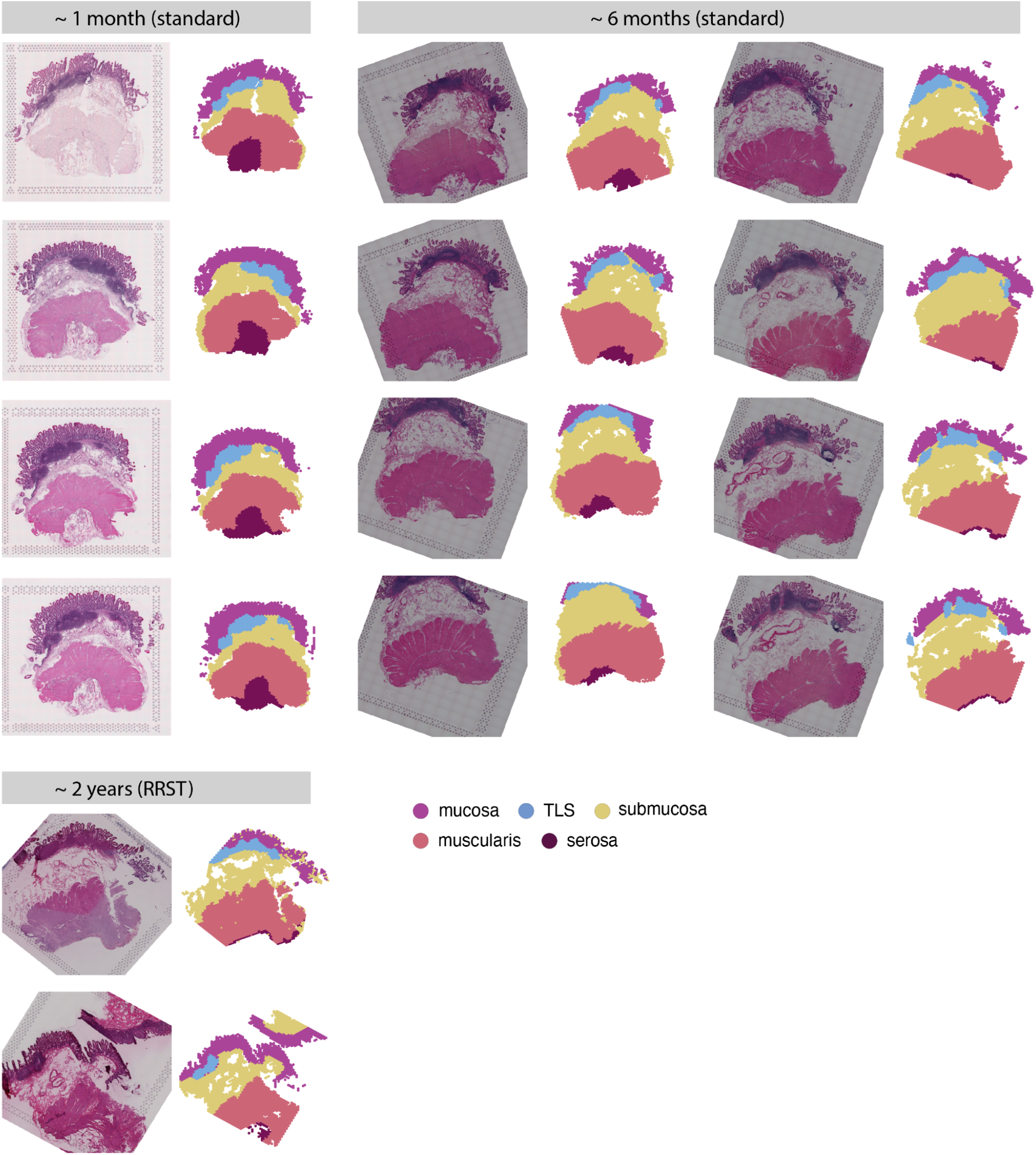
Manual annotation of small intestine data. Each of the 14 spatial transcriptomics datasets were divided into 5 regions: mucosa (intestinal epithelium), submucosa, Tertiary Lymphoid Tissue (TLS), muscularis and serosa.

**Supplementary Figure 10:**
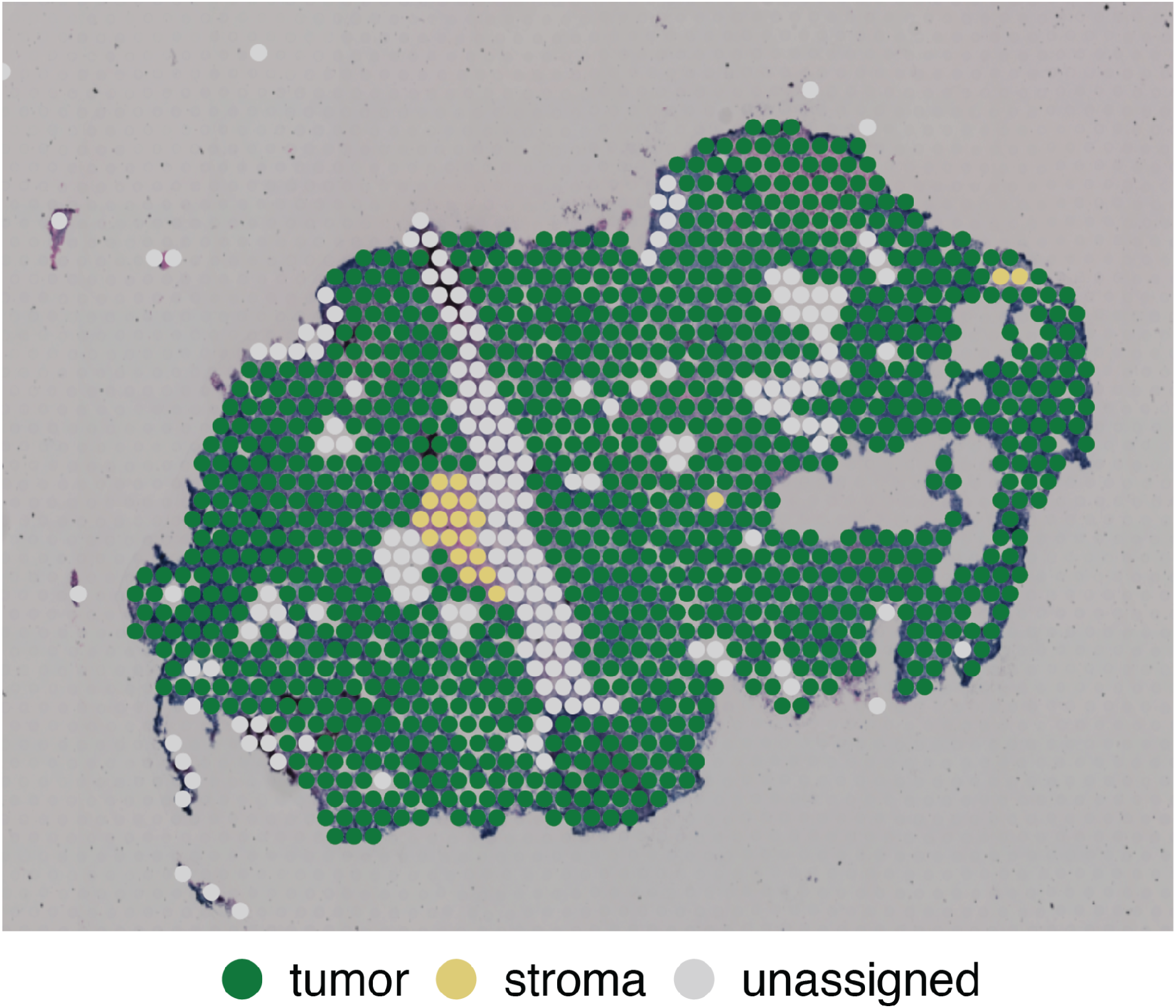
Annotations made by a pathologist of a medulloblastoma tissue section.

**Supplementary Figure 11:**
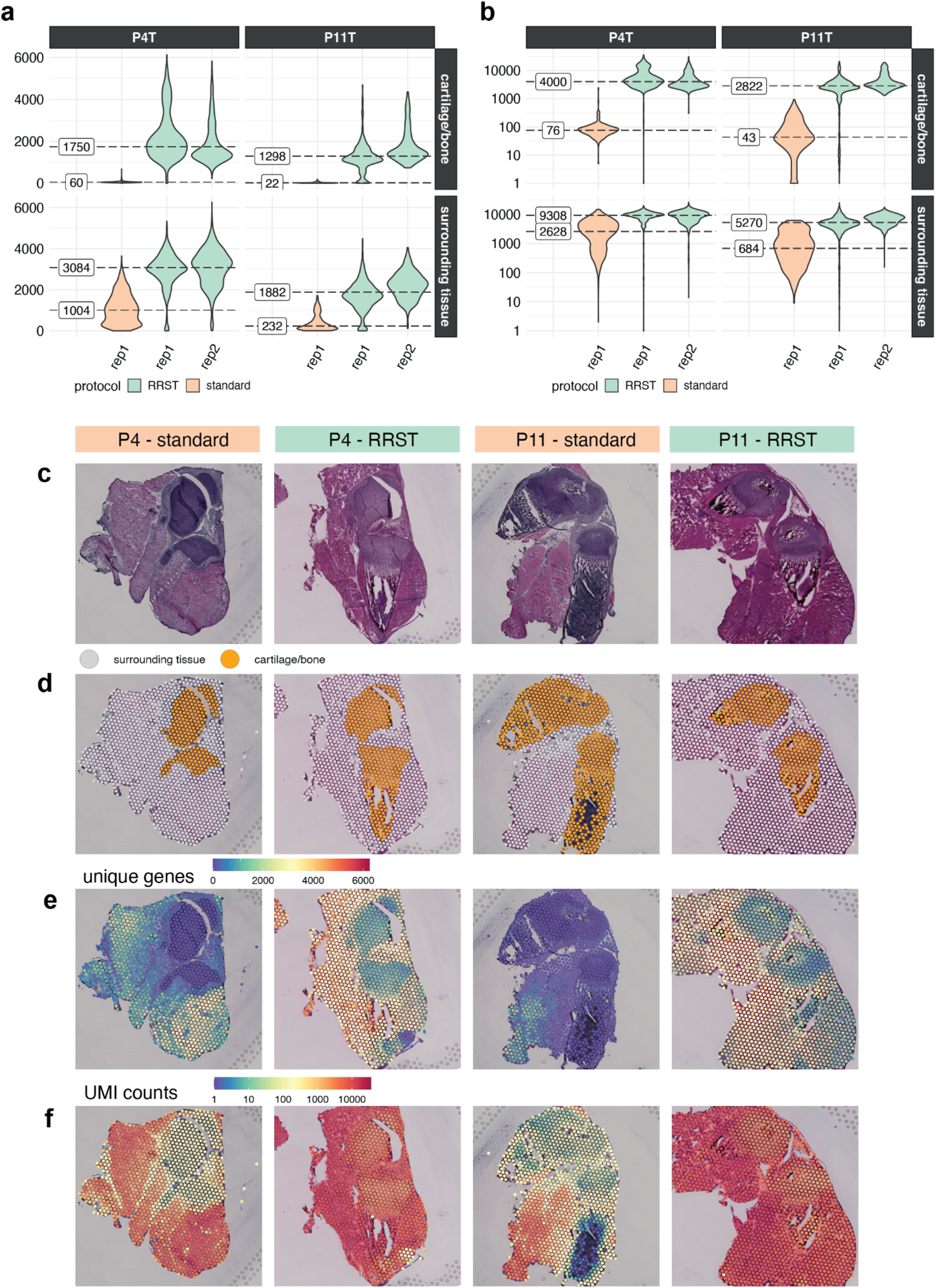
Comparison between mouse bone data obtained with standard Visium and RRST. **a)** Distribution of unique genes per spot for all tissue sections visualized as violin plots. The plots are split by age (P4T or P11T) in columns and by tissue region (cartilage/bone or surrounding tissues) in rows. The median number of unique genes is highlighted on the left side of the violin plots for each tissue section, tissue region and protocol. **b)** Distribution of UMI counts per spot for all tissue sections visualized as violin plots with the y-axis converted into log10-scale. The plots are split by age (P4T or P11T) in columns and by tissue region (cartilage/bone or surrounding tissues) in rows. The median number of UMI counts is highlighted on the left side of the violin plots for each tissue section, tissue region and protocol. **c)** H&E images for 4 representative tissue sections, one for each age group and protocol. **d)** Spots colored by tissue region: cartilage/bone or surrounding tissue. **e)** Distribution of unique genes overlaid on H&E images. **e)** Distribution of UMI counts overlaid on H&E images. The colorbar represents log10-scaled UMI counts.

**Supplementary Figure 12:**
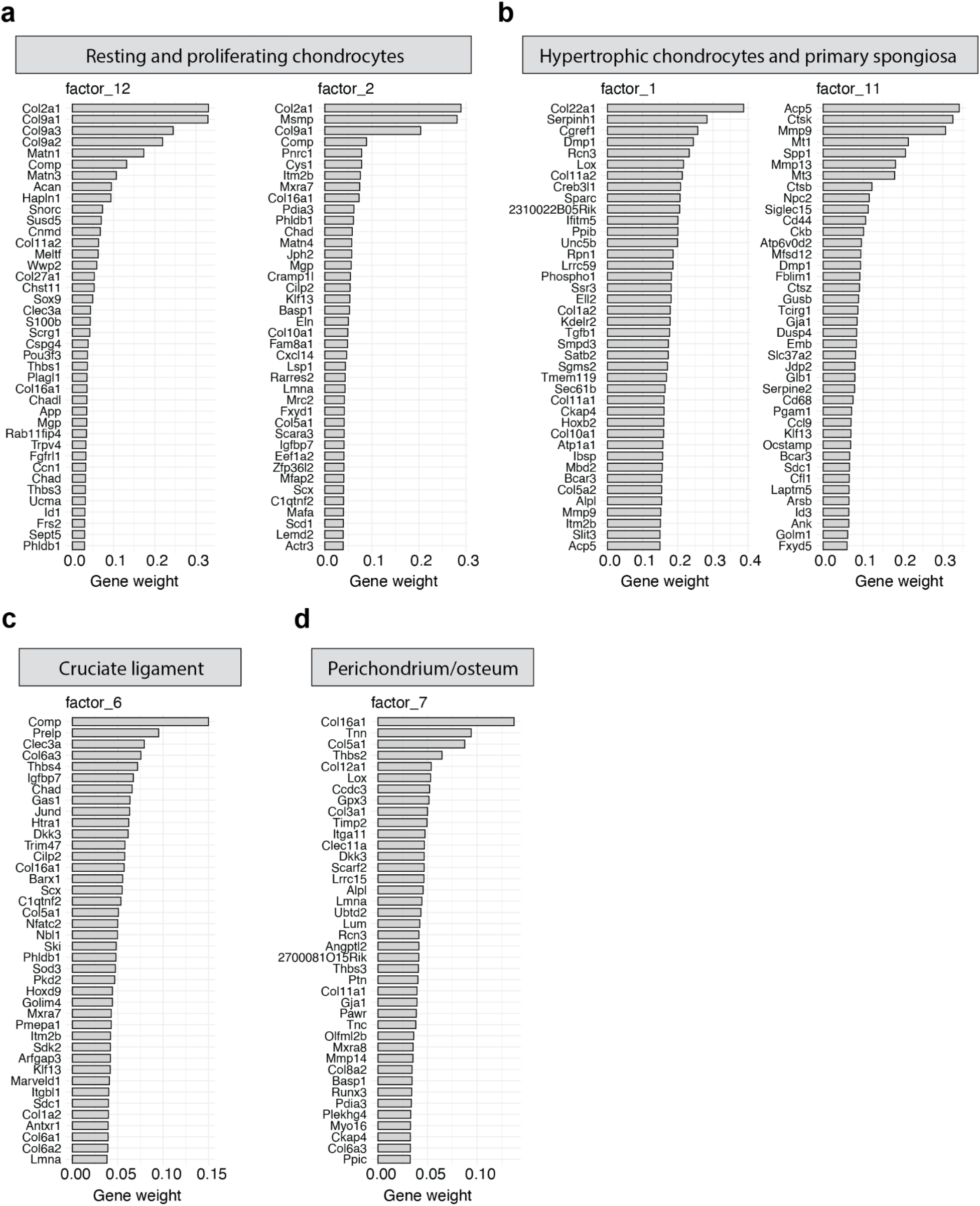
NNMF analysis identified functions containing markers of resting and proliferating chondrocytes. **a)**, hypertrophic chondrocytes and cells within the primary spongiosa **(b)**, cruciate ligament **(c)** and cells at the perichondrium/periosteum **(d)**, which appeared in the expected, discrete anatomical locations (which are shown in **Fig. 6**).

